# RUNX1-ETO induction rapidly alters chromatin landscape and growth of a specific sub-population of hESC-derived myeloid precursor cells by interfering with RUNX1 regulation

**DOI:** 10.1101/748921

**Authors:** Monica Nafria, Peter Keane, Elizabeth S. Ng, Edouard G. Stanley, Andrew G. Elefanty, Constanze Bonifer

**Author notes:** equal contribution (Monica Nafria, Peter Keane). Joint corresponding authors Constanze Bonifer, Andrew Elefanty.

## Abstract

Acute myeloid leukemia is a hematopoietic malignancy caused by recurrent mutations in genes encoding transcriptional, epigenetic and/or signaling regulators. The t(8;21) translocation generates the aberrant transcription factor RUNX1-ETO whose expression can be detected *in utero* but is insufficient to cause overt disease. Although patients harboring cells with the t(8;21) translocation have acquired additional mutations and show extensive epigenetic reprogramming, the effects directly attributable to RUNX1-ETO expression are unclear. To address this question, we used a human embryonic stem cell differentiation system capable of forming definitive human myeloid progenitor cells to express RUNX1-ETO in an inducible fashion. Induction of RUNX1-ETO causes extensive chromatin reprogramming by interfering with RUNX1 binding, blocks differentiation and arrests cellular growth, whereby growth arrest is reversible following RUNX1-ETO removal. Single cell gene expression analyses show that RUNX1-ETO induction alters the differentiation of a defined sub-population of progenitors, indicating that oncoprotein-mediated transcriptional reprogramming is highly target cell specific.

## Introduction

Hematopoietic development and differentiation are regulated through a hierarchical network of DNA-sequence-specific transcription factors. Genetic alterations affecting such regulators impair the balance of interactions within their corresponding transcriptional network, leading to a disturbance of differentiation and enhanced self-renewal. Acute Myeloid Leukemia (AML) is a heterogeneous disease marked by proliferation of neoplastic cells with impaired myeloid differentiation. The t(8;21) translocation, accounting for approximately 10% of all AML, fuses the DNA-binding domain of the hematopoietic master regulator RUNX1 to almost the entire ETO protein (Miyoshi et al., 1991). The resulting RUNX1-ETO fusion protein phenotypically functions as a dominant-negative version of RUNX1 by blocking hematopoietic development both *in vivo* and *in vitro* (Regha et al., 2015; Yergeau et al., 1997). It recruits histone deacetylase complexes to RUNX1-binding sites through its ETO moiety, resulting in repression of genes that regulate hematopoietic differentiation (Lutterbach et al., 1998; Regha et al., 2015). Experiments depleting RUNX1-ETO in established AML cells have shown that it is required to maintain leukemic growth (Ptasinska et al., 2012), but have also demonstrated that RUNX1-ETO regulated gene expression is complex with multiple genes being up- and down-regulated after knock-down (Ptasinska et al., 2014), indicating that the entire transcriptional network of such cells is rewired in presence of the fusion protein.

The t(8;21) translocation can occur early during development and has been detected *in utero* (Wiemels et al., 2010), indicating that its presence does not interfere with general hematopoietic differentiation in human embryos after formation of progenitor cells. Moreover, t(8;21) patients in remission can harbour pre-leukemic stem cells carrying the translocation but lacking secondary mutations, which may serve as a reservoir for relapse (Miyamoto et al., 2000; Shima et al., 2014). These findings agree with those found in experiments modelling the disease in mice demonstrating that RUNX1-ETO alone is not sufficient to cause AML and that secondary mutations are necessary (Higuchi et al., 2002; Yuan et al., 2001). Given that leukemia development requires the acquisition of multiple genetic aberrations, the study of primary cells from patient leukemic samples does not allow easy discrimination of the epigenetic impact of RUNX1-ETO alone. A number of studies examined the development of AML using inducible RUNX1-ETO expression in mice or constitutive expression in human cells *in vitro*. However, these experiments suffered from the fact that AML development was slow (Cabezas-Wallscheid et al., 2013), the proliferation of cells in the culture dish implied the requirement for a selection step (Mandoli et al., 2016; Mulloy et al., 2002, 2003), RUNX-ETO was induced at higher levels than seen in patients (Mulloy et al., 2002, 2003), or assays were performed in established cell lines harbouring additional mutations (Martens et al., 2012; Ptasinska et al., 2012, 2014), or in *in vitro* differentiated yolk sac-like progenitors that are unlikely to represent the proper target cell types (Mandoli et al., 2016). These caveats hinder the understanding of the earliest, unperturbed epigenetic reprogramming events occurring in a human setting after RUNX1-ETO induction. For this reason, we genetically altered human pluripotent stem cells to inducibly express RUNX1-ETO in response to doxycycline, and used an *in vitro* system of hematopoietic differentiation that biases cultures towards definitive multipotent hematopoietic progenitor cells (Ng et al., 2016).

Our experiments showed that high levels of RUNX1-ETO had a detrimental effect on hematopoiesis. However, levels of *RUNX1-ETO* expression that matched expression of RUNX1 in immature clonogenic blood progenitors were compatible with cell survival. Within 24-hours of *RUNX-ETO* induction, cells became quiescent, downregulated hematopoietic differentiation, cell cycle and DNA repair genes, but up regulated MAPK and VEGF signalling pathway genes. In contrast to un-induced cells, these cells could survive for months in vitro without proliferating. Strikingly, an enhanced clonogenic capacity of these immature, quiescent RUNX1-ETO expressing cells was only revealed following the removal of doxycycline, ceasing RUNX1-ETO transcription and allowing cells to proliferate and differentiate. Chromatin immunoprecipitation and chromatin accessibility assays showed that RUNX1-ETO binding led to wide-spread loss of chromatin accessibility at RUNX1 binding sites and substantially altered the RUNX1-controlled transcriptional program. Single cell RNA-Seq experiments demonstrated that RUNX1-ETO induction exerted its main impact on a defined early precursor population by down-regulating the myeloid master regulator PU.1 but had little effect at later stages or different lineages. Our study therefore sheds light on the earliest events occurring in t(8;21) cells after the first leukemic mutation in a human primary cell setting and demonstrates a dissociation between the block in differentiation and cell proliferation, considered to be the two hallmarks of leukemia.

## Results

### Expression of RUNX1-ETO leads to a reversible differentiation and growth arrest of human early hematopoietic progenitor cells

In order to analyse the effects of RUNX1-ETO induction in defined cell types, we generated inducible RUNX1-ETO human ESC lines. The parental line used was a previously generated human H9 ESC dual reporter cell line (denoted SOX17^mCHERRY/w^RUNX1C^GFP/w^) carrying an *mCHERRY* gene in the *SOX17* locus, marking arterial endothelium (Clarke et al., 2013), and a *GFP* gene in the *RUNX1* locus, resulting in expression of GFP from the distal promoter (*RUNX1C*) and hence marking hematopoietic stem and progenitor cells (Ng et al., 2016) (Figure S1A). *RUNX1C* is the dominant *RUNX1* isoform in fetal liver blood progenitors (Sroczynska et al., 2009). In contrast to *RUNX1B*, *RUNX1C* expression is restricted to hematopoietic cells and defines the subset of CD34+ cells with clonogenic and bone marrow homing activity (Ng et al., 2016). This strategy allowed us to track the progression of *in vitro* cell differentiation (Figures 1A, B, and S1B), facilitating the distinction between endothelial and hematopoietic cells. It also allowed us to monitor the endothelial-hematopoietic transition (EHT), the process by which hematopoietic progenitor cells form and detach from the endothelium which is accompanied by a switch in *RUNX1* promoter usage (Bertrand et al., 2010; Boisset et al., 2010; De Bruijn et al., 2002; Jaffredo et al., 1998; Kissa and Herbomel, 2010; Lam et al., 2010; North et al., 2002). We derived definitive hematopoietic progenitors from ESC lines using a protocol that included SB431542 and CHIR (an ACTIVIN inhibitor and WNT agonist, respectively) from day 2-4 to pattern cells towards an intra-embryonic, definitive *HOXA+* fate (Ng et al., 2016) (Fig 1A). By differentiating the dual reporter cell line using this protocol, we were able to visualize RUNX1C+ progenitors emerging from SOX17+ hemogenic endothelium and forming cell clusters, resembling progenitor formation within the week 5 human aorta-gonad-mesonephros (AGM) (Ng et al., 2016) (Fig 1B, S1B).

**FIGURE 1.**
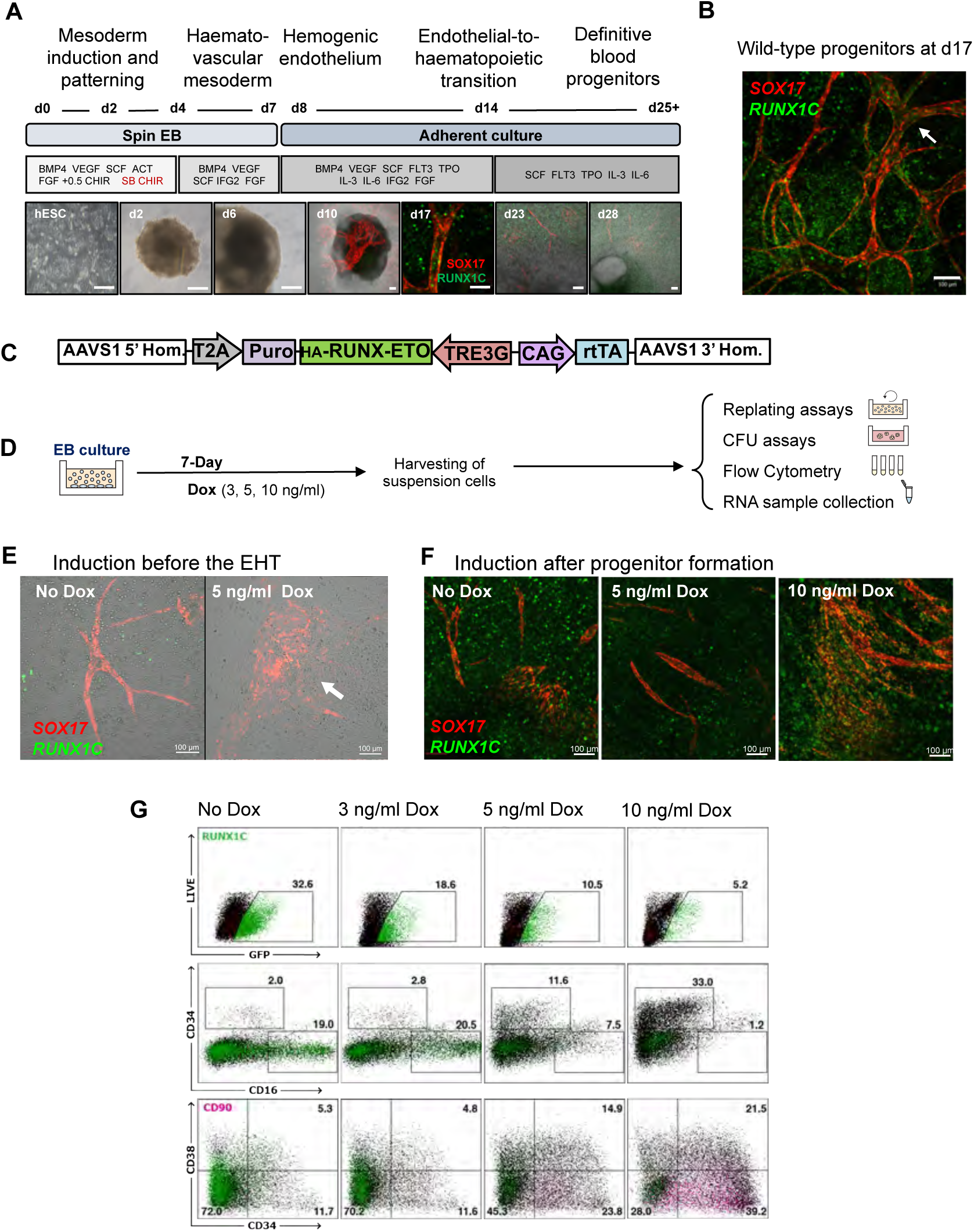
Expression of *RUNX1-ETO* leads to a reversible differentiation and growth arrest of human early hematopoietic progenitor cells. A) Protocol for *in vitro* human definitive hematopoietic differentiation as spin embryoid bodies (EBs). Developmental stages, time course and growth factors used are indicated. Epi-fluorescence (hESC, d2 and d6) and confocal images (d10-d28) representative of each stage are shown. A pulse of SB and CHIR (red) was included from d2-4. Human Embryonic Stem cells (hESC) are shown at ∼50% confluency on a feeder layer. EBs (d2, d6) appear as opaque round structures, and are surrounded by adherent stroma, endothelium and blood cells from d10. Fluorescence and brightfield channels are merged in images corresponding to d10, d13 and d28. Scale bar: 100 μm. SOX17 (mCHERRY, red) expression marks vascular structures and RUNX1C (GFP, green) marks hematopoietic progenitors. B) Confocal image of a differentiation culture at d17 showing RUNX1C+ progenitors being generated from SOX17+ vascular structures of endothelial cells, resembling embryonic AGM hematopoiesis. SOX17 (CHERRY; red), RUNX1C (GFP; green). Arrow points groups of progenitor cells resembling the embryonic intra-aortic hematopoietic cell clusters. Scale bar: 100 μm. C) Schematic representation of the transgene targeting into the AAVS1 locus. Knock-in was performed via TALEN-mediated homologous recombination into the AAVS1 locus of the SOX17^mCHERRY/w^ RUNX1C^GFP/w^ human embryonic stem cell line H9. The integrated sequence includes an HA-tagged *RUNX1-ETO* cDNA under control of a tetracycline-inducible expression system (TRE-3G), the reverse tetracycline activator (rtTA) controlled by a chicken beta-actin promoter (CAG) and a puromycin resistance gene with an upstream T2A sequence to link its expression to the AAVSI gene. D) Experimental strategy for the evaluation of RUNX-ETO induced by Dox (0, 3, 5 or 10 ng/ml). The non-adherent hematopoietic cell fraction was harvested 7 days after induction and used flow cytometry analysis, colony forming unit and replating assays. E) Confocal images of hematopoietic cultures showing the disruptive effects of RUNX1-ETO induction before the EHT on vasculogenesis and blood formation. Images are representative of uninduced and cultures induced at d10 (5 ng/ml Dox) for 7-days. SOX17 (CHERRY) and RUNX1C (GFP). Fluorescence and brightfield channels are merged. Scale bar: 100 μm. F) Confocal images of hematopoietic cultures showing the effect of RUNX1-ETO on formation of blood progenitors. *RUNX1-ETO* expression at an equivalent level to that of endogenous RUNX1 (5 ng/ml Dox) allows vasculogenesis and blood formation to occur, whilst higher *RUNX1-ETO* expression levels (10 ng/ml Dox) result in formation of abnormal vascular structures and reduced blood formation. Images are a representative of uninduced and cultures induced at d21 with 5 or 10 ng/ml Dox for 7-days. SOX17 (CHERRY) and RUNX1C (GFP). Scale bar: 100 μm. G) *RUNX1-ETO*-expressing cultures retain markers of immature myeloid progenitors. Flow cytometry analysis of the floating fraction of d34 hematopoietic progenitors upon 7-day RUNX1-ETO induction. Results are a representative of three biological replicates with comparable results with induction at different time points after the EHT.

We next generated cell lines containing an N-terminal hemagglutinin (HA)-tagged *RUNX1-ETO* cDNA, expressed in a doxycycline (Dox)-inducible manner, targeted to the safe harbour AAVS1 locus (Fig 1C). Induction of HA-*RUNX1-ETO* occurred in the vast majority of cells (Fig S1C, S1D). We first determined the Dox concentration required to induce *RUNX1-ETO* expression to levels in hematopoietic progenitors that mimicked the balanced ratio of RUNX1-ETO:RUNX1 expression observed in AML patients (Fig S1E), concluding that 3 - 5 ng/ml was the most appropriate concentration. Higher Dox concentrations (100-500 ng/ml) that further increased RUNX1-ETO levels completely abrogated blood formation (data not shown). To examine the dose response of hematopoiesis to RUNX1-ETO expression we induced differentiating cells with either 3, 5 or 10 ng/ml Dox (Fig 1D). Dox induction before the endothelial to hematopoietic transition (EHT) resulted in severe disorganization of the SOX17+ vasculature and an overall reduction of blood cells that also appeared phenotypically abnormal (Fig 1E, S1F). Abnormal progenitors either lacked GFP (RUNX1C) expression, failed to downregulate SOX17 (CHERRY+) or co-expressed both SOX17 and RUNX1C (CHERRY and GFP positive). However, induction after the EHT allowed the generation of phenotypically normal blood cells, with only 10 ng/ml Dox reducing blood cells numbers (Fig 1G). Expression of RUNX1-ETO also affected the nature of the hematopoietic cells present in our cultures causing a Dox-dependent decrease of RUNX1C+ and CD16+ myeloid cells and an increase of CD34+CD38-CD90+ populations resembling immature blood progenitors (Fig 1G).

RUNX1-ETO induction reduced colony forming ability in CFU assays in Dox-containing methylcellulose medium in a dose-dependent manner (Fig 2A, S1G left panel). However, CFU activity was restored once Dox was removed from the methylcellulose (Fig 2A, S1G right panel), indicating that reversibility of the RUNX1-ETO-dependent proliferation block. The presence or absence of Dox did not impact on colony size or morphology, suggesting that the clonogenic cells differentiated normally following removal of Dox (data not shown). Moreover, prolonged culture of hematopoietic cells in the presence of low levels of RUNX1-ETO (3 - 5 ng/ml doxycycline) prolonged cell survival without cell proliferation (Fig 2B,C; Fig S1H, I). In addition, presence of RUNX1-ETO caused a profound decrease in DNA-synthesis activity due to an arrest in the G1 phase of the cell cycle, as measured by BrDU incorporation, without an increase in cell death (Fig S1J).

**Figure 2.**
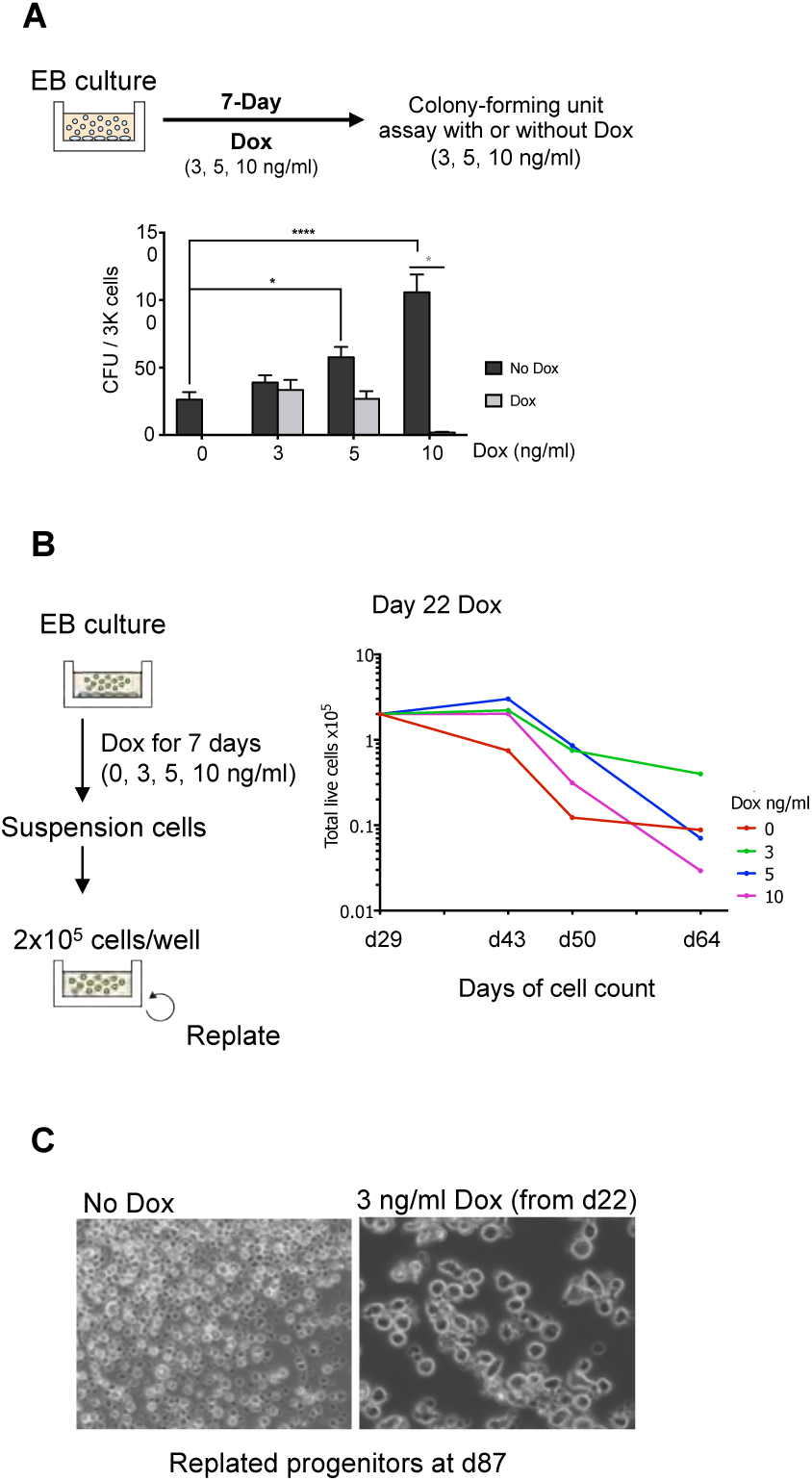
A) RUNX1-ETO induction causes a reversible block in colony forming ability. Top panel; Diagram depicting the experimental strategy. EB cultures were treated with 3, 5, or 10 ng/ml Dox for 7 days and suspension cells were plated in methylcellulose for colony-forming unit (CFU) assays in either presence or absence of Dox. Below; CFU assay of progenitors from treated EB cultures at day 24 for 7 Days, plated in methylcellulose in the presence (light grey) or absence (dark grey) of Dox. Data are from three independent biologic replicates using two different clones. CFU assays conducted in triplicates with 3,000 cells plated per well. Grey-coloured (*): Multiple T-test, Statistical significance determined using the Holm-Sidak method, with alpha = 0.05. Each row was analyzed individually, without assuming a consistent SD. Black-coloured (*): Two-way ANOVA, Statistical significance determined using the Dunnett’s multiple comparisons test. B) Induction of RUNX1-ETO at low levels (3 ng/ml, 5 ng/ml Dox) enhances the survival of a subset of progenitor cells as compared to un-induced cells, without increasing proliferative capacity. Left; Schematic diagram of the replating assays. EB cultures were treated at different stages of hematopoietic differentiation with 0, 3, 5 or 10 ng/ml Dox for 7 days. Floating progenitors were plated on Matrigel-coated wells at 2 x10^5^ cells/well in the corresponding Dox concentration and were serially passaged each week. Right; Cell count upon replating assays of hematopoietic progenitors from cultures treated at d22 with Dox, showing one representative of three biological replicates. Cell growth was measured at the indicated times. C) RUNX1-ETO expressed at low levels allows survival of cells until day 87. Brightfield images of hematopoietic progenitors from replating assays at d87 of differentiation that are uninduced (left) or treated with 3 ng/ml Dox from d22 onwards (right). Images are taken using the same magnification.

In summary, our experiments show that the expression of balanced levels of *RUNX1-ETO* in developing human blood progenitor cells leads to a reversible block in differentiation with growth arrest and prolongation of cell survival.

### RUNX1-ETO induction leads to cell-type and dose-dependent changes in gene expression

We previously demonstrated that RUNX1-ETO reshapes the transcriptional network in a developmental stage-specific manner and that the outcome is dependent on the cell type in which the oncogenic event occurs (Regha et al., 2015). To examine the molecular basis of this finding in human cells, we examined genes induced by RUNX1-ETO in two progenitor cell populations (SOX17^−^ CD45^+^CD34^+^RUNX1C^+(GFP+)^) and SOX17^−^CD45^+^CD34^+^RUNX1C^−(GFP−)^) isolated from d22 differentiating cultures with or without prior induction of RUNX-ETO by 24 hours of doxycycline treatment (Fig S2A). Both populations comprised CD34+CD45+ progenitors but the molecular distinction between RUNX1C+ and RUNX1C-populations was not known. Analysis of differential gene expression between RUNX1C+ and RUNX1-cells showed over 500 and 600 genes up- and down-regulated, respectively, demonstrating an intrinsically different nature of the two cell populations (Fig S2B). Expression levels of *RUNX1* and *SPI1* were similar but RUNX1C-cells expressed high levels of monocyte-specific genes such as *IRF8, CSF1R* and *CD14* indicating the presence of maturing myelomonocytic cells (Supplemental dataset 2). RUNX1C+ cells expressed higher levels of *MYB, GATA2* and *GFI1* as well as the erythroid regulators *GATA1* and *KLF1.* After induction of RUNX1-ETO with 5 ng/ml Dox, both RUNX1C- and RUNX1C+ cell populations up- and down-regulated similar number of genes (Fig S2C). The actual RUNX1-ETO responsive gene expression program was also different (Fig 3A, Supplementary Dataset 3). In accordance, up- and down-regulated KEGG pathways in gene sets from cells responding to 5 ng/ml or 10 ng/ml Dox induction differed as well (Fig 3B, Supplementary Dataset 4). RUNX1C+ cells down-regulated cell cycle and DNA-replication genes (such as *BRCA1*, *BUB1B* and *RNASEH2B*) and up-regulated a large number of signalling genes (such as *MAPK3* and *JUN*), whilst RUNX1C-cells down-regulated genes belonging to hematopoietic lineage pathways (such as *CEBPA*, *IL4* and *KIT*) and up-regulated only a subset of the genes upregulated in the RUNX1C+ population. Moreover, the gene expression response to RUNX1-ETO induction in RUNX1C+ cells was highly dose-dependent (Fig 3C, S2D). Furthermore, RUNX1-ETO induction yielded highly heterogeneous changes in gene expression with distinct subsets of genes responding differently to the oncogene dosage (Fig S2E). For example, expression of the stem cell regulator *GATA2* and the *WT1* gene, decreased, along with cell cycle *genes* (*BUB1, CHEK2, CCNB1)* and the growth factor receptor gene *KIT* (Fig 3D). In contrast, genes involved in signalling pathways (*MAPK3)* and immediate early response genes (*FOS, JUN)* were up-regulated (Fig 3D). These gene expression data were concordant with the observed cell cycle arrest and demonstrate that RUNX1-ETO impacts on distinct cell types in a different fashion, highlighting the importance of inducing RUNX1-ETO in the appropriate cell type for the understanding of the leukemic phenotype.

**FIGURE 3:**
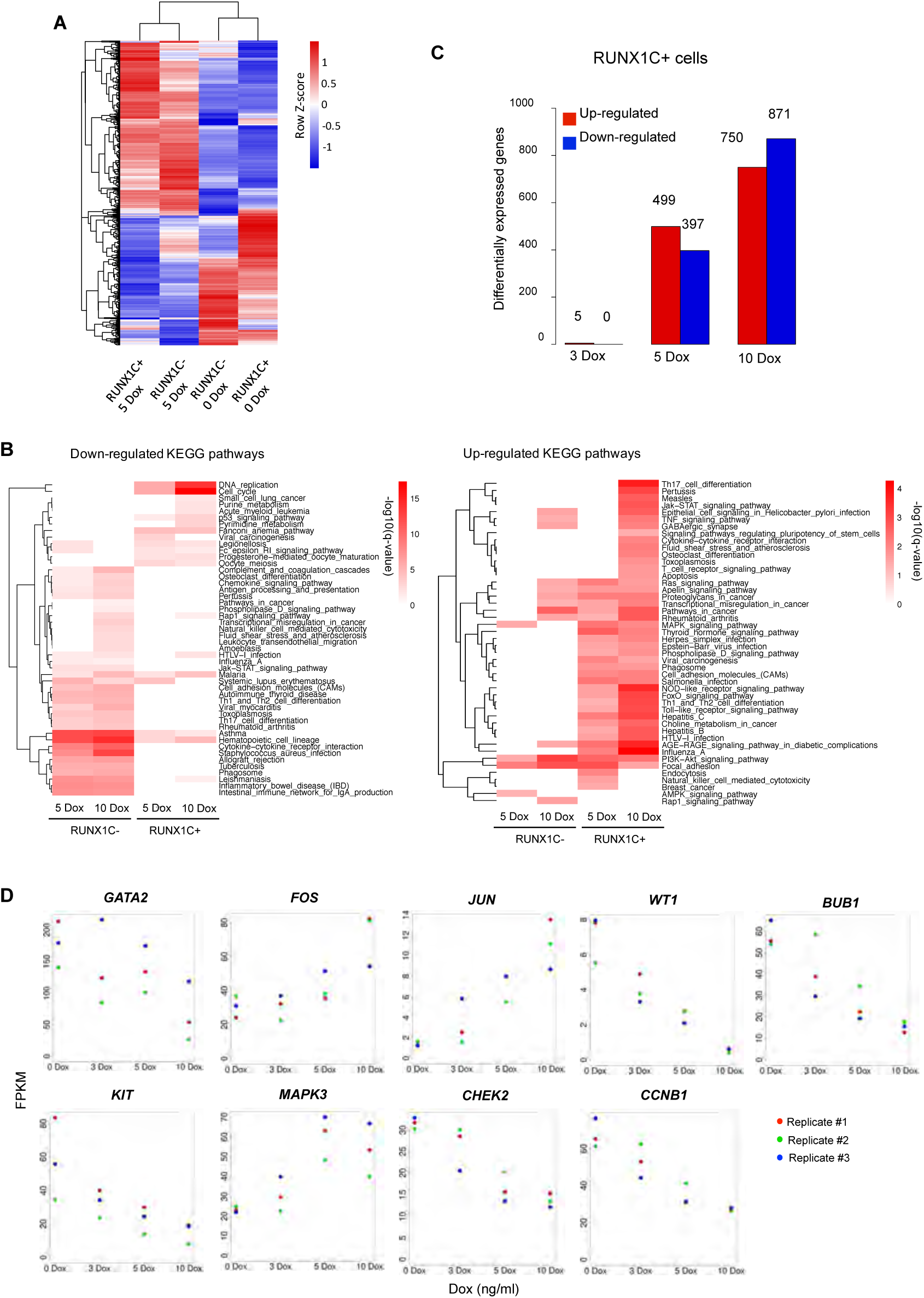
RUNX1-ETO induction leads to cell-type and dose-dependent changes in gene expression. A) Clustering of gene expression data for RUNX1C- and RUNX1C+ (CD45+ CD34+) sorted cells both wild-type and after 24-hour RUNX1-ETO induction using 5 ng/ml dox. The figure includes all genes which showed up/down regulation after RUNX1-ETO induction in either the RUNX1C+ or RUNX1C-cell populations. B) Heatmap after KEGG pathway analysis depicting clustering of differentially expressed genes associated with KEGG terms upon RUNX1-ETO induction (5 ng/ml Dox and 10 ng/ml Dox) in both RUNX1C- and RUNX1C+ sorted populations. Red colour intensity reflects the enrichment significance of the terms in −log10 (q value). C) Bar graph depicting differentially expressed genes between No Dox and Dox-treated (3, 5 and 10 ng/ml) CD45+ CD34+ RUNX1C+ sorted cells. D) Examples of individual genes differentially regulated in CD34+ RUNX1C+ progenitors in response to RUNX1-ETO induction (3, 5 10 ng/ml Dox) for 24h. N=3. Each color represents a distinct biological replicate.

### RUNX1-ETO induction causes extensive global chromatin reorganisation and disrupts the binding of RUNX1 at distal elements

In order to understand the RUNX1-ETO-mediated chromatin reprogramming and changes in transcription factor binding in RUNX1C+ cells, we next analysed open chromatin regions using the Assay for Transposase-Accessible Chromatin with high-throughput sequencing (ATAC-Seq) and protein binding by performing Chromatin Immunoprecipitation (ChIP) experiments (Fig 4A). Induction of RUNX1-ETO dramatically shifted the accessible chromatin landscape (Fig 4B (ATAC-Seq)), an example for this being the *RASSF5* locus shown in Fig 3C. A large number of accessible sites were lost (5419) and gained (4112) within 24h of Dox induction. Lost sites were associated with down-regulated gene expression (Fig 4B (gene expression)) and were enriched in binding motifs for important members of hematopoietic transcription factor families, such as PU.1 (but not ETS) as well as GATA, RUNX and C/EBP family members (Fig 4B (motif density plots) and Fig 4D). These results were confirmed by ChIP-Seq experiments showing loss of RUNX1 binding across all RUNX1-ETO bound sites as well as a reduction of the active histone marks H3K27 acetylation (Ac) and H3K4 trimethylation (me3) (Fig 4B, Fig 4E). These losses were most pronounced on distal elements (Fig S3 C, D) whereas promoters were less affected (Fig S3 A, B).

**FIGURE 4:**
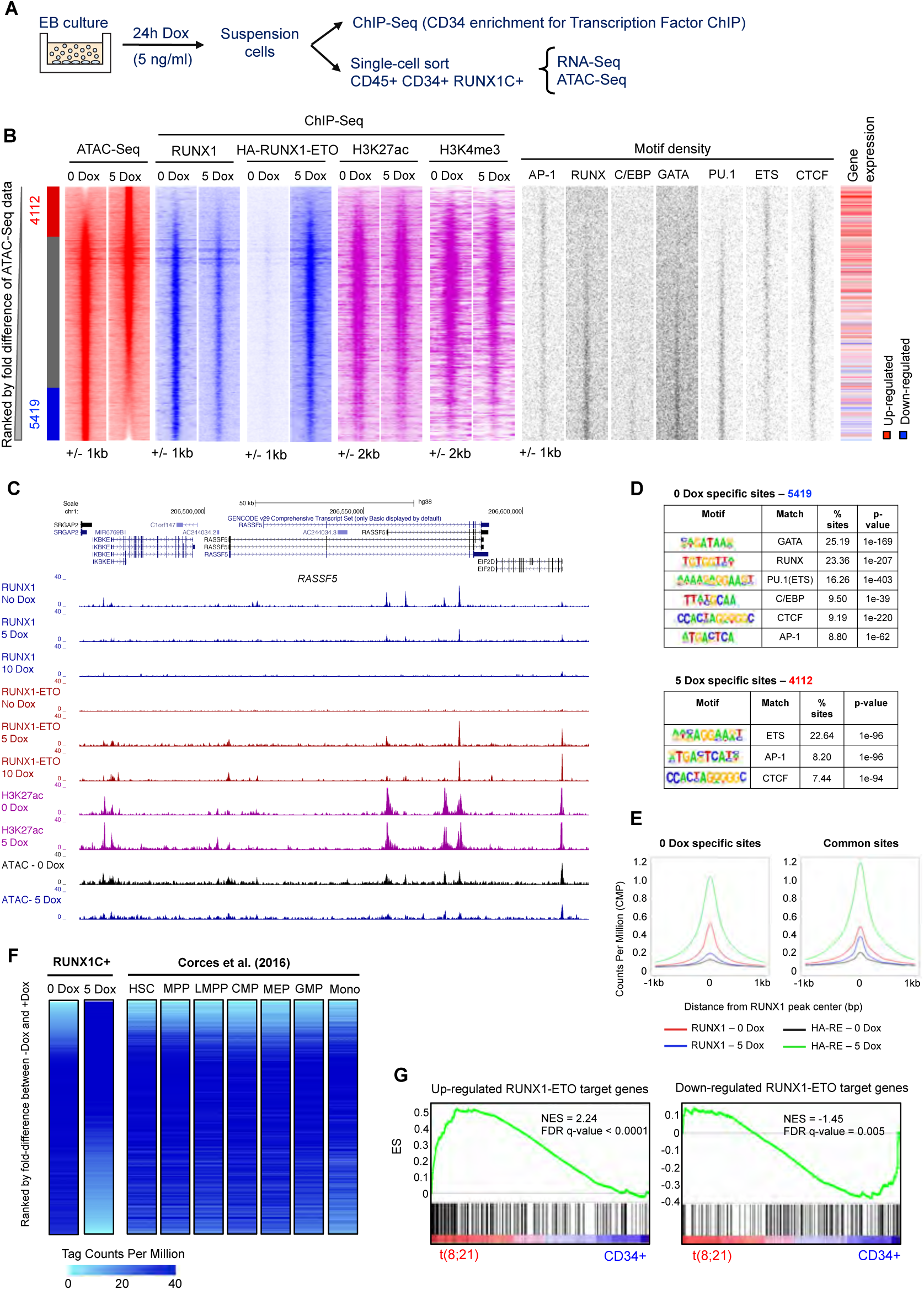
RUNX1-ETO induction causes extensive global chromatin reorganisation and blocks the binding of RUNX1. A) Sorting strategy and downstream analyses after 24h Dox induction of RUNX1-ETO. B) Heatmaps depicting accessible chromatin regions ranked by fold difference between the 0 and 5 Dox RUNX1C+ treated samples. ATAC-Seq peaks were considered sample-specific when displaying a greater than 2-fold enrichment compared to the other sample. Sample-specific sites and number of peaks are indicated alongside, being: red the 5-Dox specific, blue the 0-Dox specific and grey the shared peaks. ChIP-Seq enrichment for RUNX1, HA-RUNX1-ETO, H3K27ac and H3K4me3 in each sample, motif density plots and gene expression at these sites are ranked along the same coordinates as the ATAC-Seq peaks. C) Genome browser screenshot depicting RUNX1, HA-RUNX1-ETO, H3K27ac ChIP-Seq and ATAC-Seq tracks for the indicated samples at the *RASSF5* gene locus. D) Motif enrichment analysis in 0 and 5 Dox-specific peaks. E) Average profiles for RUNX1 and RUNX-ETO ChIP-Seq data centred on RUNX1 binding peaks (+/− 1000bp from peak centre) in the 0 and 5 Dox-specific peaks. F) Comparison of chromatin accessibility in RUNX1C+ cells (0 and 5 Dox-treated samples) to myeloid progenitor cell types from Corces et al (2016). Heatmaps show ATAC-Seq tag-counts ranked by fold difference between the 0 and 5 Dox treated RUNX1C+ samples. ATAC-Seq tag counts from distinct myeloid progenitor cell types (Corces et al., 2016) are ranked along the same coordinates as the 0 Dox ATAC-Seq peaks. Colour intensity reflects tag counts per million, with red representing closed chromatin. G) RUNX1-ETO-induced human ESC-derived progenitors share the same gene expression profile as in t(8;21) AML patients. Gene Set Enrichment Analysis (GSEA) correlating upregulated (upper panel) and downregulated (lower panel) RUNX1-ETO target genes between CD45+CD34+RUNX1C+ cells following 24-hour RUNX1-ETO induction (5 ng/ml Dox) and the gene expression profile of the RUNX1-ETO targets in t(8;21) patients. ES: Enrichment Score, NES: Normalized Enrichment Score, FDR: False discovery rate.

The binding pattern of RUNX1-ETO showed a large overlap with that of RUNX1 and the majority of genes bound by both factors lost RUNX1 binding after RUNX1-ETO induction (Fig S3E, green circle). Around 40% of both upregulated and downregulated genes upon induction were RUNX-ETO target genes (Figure S3F). Moreover, approximately half of the downregulated and upregulated RUNX-ETO target genes were also RUNX1 targets that would lose RUNX1 binding upon induction (Supplemental Dataset 5). However, a small proportion of the differentially expressed RUNX1 targets that lose binding upon induction, do not appear to be targeted by RUNX1-ETO (Figure S3F, light colours). Taken together, our data demonstrate that induction of RUNX1-ETO dramatically alters the RUNX1 driven gene expression program, mostly by direct, but also by indirect means.

Human ESC-derived multipotent progenitor cells are transcriptionally similar to the first preHSCs developing from the human AGM (Ng et al., 2016). However, changes in the chromatin structure often precede the onset of transcription (Bonifer and Cockerill, 2017). We therefore wished to determine to what extent the chromatin landscape in our human ESC-derived progenitors resembled that of normal blood stem and progenitor cells. To this end, we compared our ATAC-Seq data derived from un-induced and induced cells to those generated from highly purified human hematopoietic precursor populations as well as monocytic cells (Corces et al., 2016) (Fig 4F). This analysis shows a strong resemblance of the overall ATAC-Seq pattern of human ESC-derived RUNX1C+ progenitors to HSC and MPP populations, but not to monocytes. RUNX1-ETO induction completely shifted this pattern, resulting in loss of open chromatin regions specific for early progenitors and appearance of new accessible sites. In human ESC-derived progenitors, RUNX1-ETO induction deregulated a similar set of genes as that found in t(8;21) AML patients when compared to normal CD34+ stem/progenitor cells (Fig 4G). This result argues that the initial RUNX-ETO mutation may account for a large portion of the altered transcriptional network in leukemia patient samples.

### RUNX1-ETO expression blocks the differentiation of a specific cell population

To identify the cells most responsive to RUNX1-ETO induction, we performed single cell (sc) RNA-Seq experiments, comparing purified CD45+CD34+RUNX1C+ cells with or without 24-hour treatment with 5 ng/ml Dox, assigning cell populations based on expression of known lineage marker genes (Fig 5A and S4A-C, Supplemental dataset 6) The clustering analysis in Figure 5B shows that the un-induced population consisted of early erythroid precursors, together with maturing erythroid and myeloid cells, and a population of less differentiated cells resembling stem and progenitor cells (Fig 5B, left panel). Overnight induction of RUNX1-ETO led to the emergence of an enriched population (Fig 5B right panel (green), S4D), herein referred as 5-Dox enriched population. Analysis of cell cycle regulated genes within this population showed a strong arrest in the G1 phase of the cell cycle (Fig 5C). Some degree of cell cycle arrest was also observed in the stem/progenitor, eosinophil and GMP-like populations (orange / purple / dark blue) but not in the erythroid (pink / red) populations. A similar result was also seen when we measured the number of genes responding to induction (Fig S4E), with the population with highest number of changing genes being the 5-Dox enriched new cell population (green), followed by the stem/progenitor population (orange) and lastly the erythroid lineage (pink and red), which showed only few responsive genes. We next projected the expression of important hematopoietic regulator genes on the different cell clusters (Fig 5D). This analysis demonstrated again the cell-type specificity of the response to RUNX1-ETO induction, since genes were differently regulated on the distinct cell clusters. The quantification of expression of individual genes (Fig S5) in the different clusters showed that the most dramatic response was the loss of *SPI1* (PU.1) expression in the 5-Dox enriched population, as well as strongly upregulated expression of the *SOX4* transcription factor gene. We also observed a reduction of *GATA2*, *CEBPA* and *GFI1B* expression. Interestingly, cells in the 5-Dox enriched population expressed the chemokine gene *CCL5*, which has roles in proliferation, metastasis and in creating an immunosuppressive environment. As shown above, transcription factors relevant for erythroid development and erythroid genes were largely unaffected, as were other genes such as *MEIS1.* Gene expression and KEGG pathway analysis of up-and down regulated genes in the 5-Dox enriched population are consistent with the results obtained from the bulk RNA-Seq data (Fig S6A,B, Supplemental Dataset 7). These results show that single cell analysis of RUNX1-ETO transcriptional dysregulation yielded similar results to the differential gene expression observed in the bulk progenitor population upon 5 Dox treatment, confirming the RUNX1-ETO-dependent down-regulation of cell cycle, replication, and interestingly, spliceosome and ribosomal genes.

**FIGURE 5:**
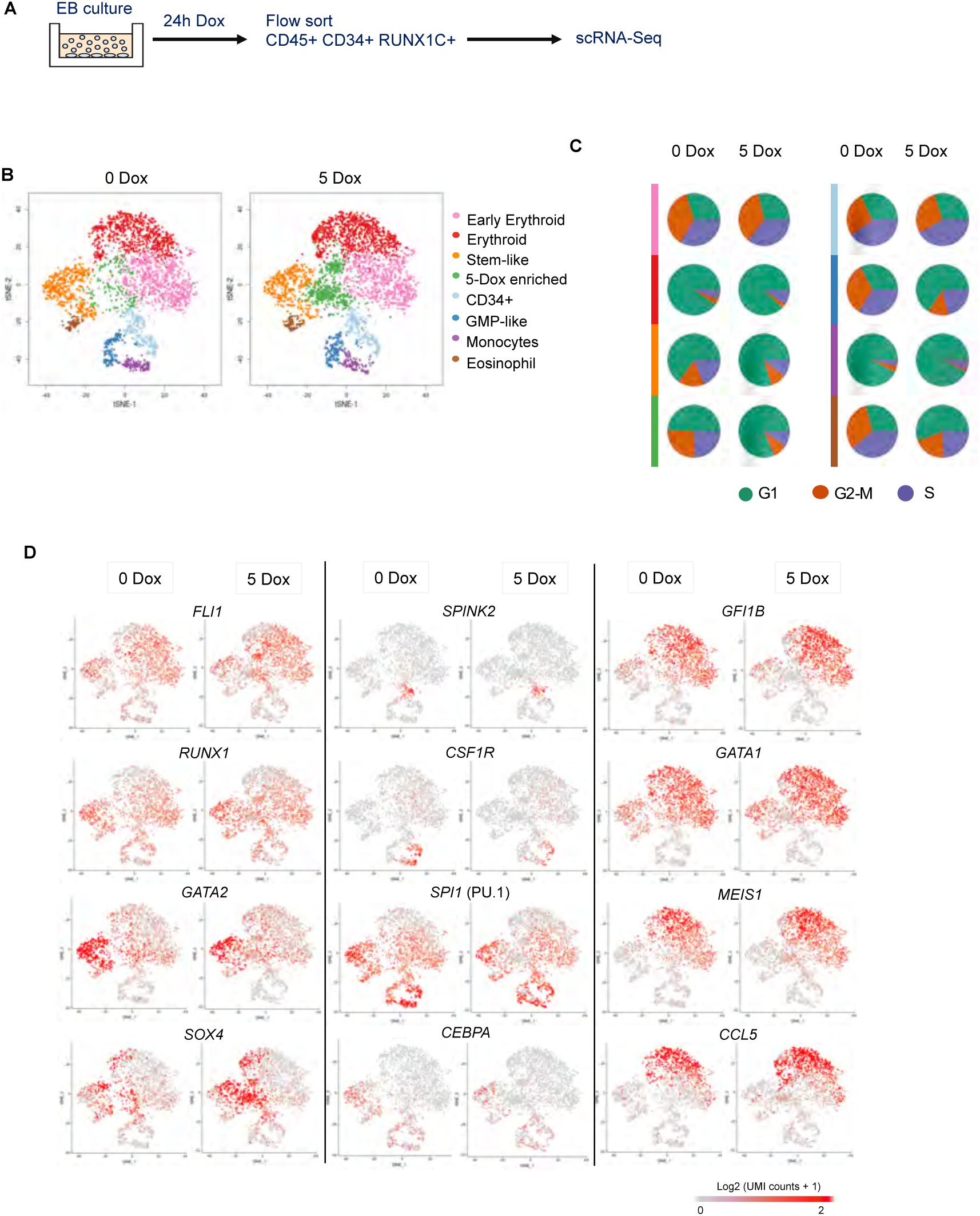
Induction of RUNX1-ETO in the CD45+CD34+RUNX1C+ population results in the emergence of a new cell population. A) Diagram of the sorting strategy for single cell (sc) RNA-Seq performed following 24h Dox induction of RUNX1-ETO. B) Two-dimensional t-SNE maps displaying 3,321 (left) and 3,814 (right) CD45+ CD34+ RUNX1C+ sorted single cells following 0 and 5 ng/ml Dox treatment, respectively. Colours represent the different clusters identified after RaceID analysis. C) Pie charts displaying the proportion of cells in each cell cycle phase (G1, G2-M and S) within each cluster as identified by expression of cell cycle regulated genes such as histone genes D) Expression of individual marker genes projected on the t-SNE maps of both untreated (0 Dox) and treated (5 Dox; 5ng/ml for 24h) scRNA populations. Colour intensity represents number of transcripts sequenced in Log2 of unique molecular identifier (UMI) counts +1.

To gain more insight into the position of RUNX1-ETO-responsive cell population within the differentiation trajectory, we performed a pseudotime (nearest neighbour) analysis (Fig 6). Un-induced cells showed a clear distribution of the different populations with the stem-like (orange) population at the apex of the hierarchy from which the erythroid and myeloid lineages branched (Fig 6A, left panel). Induction of RUNX1-ETO distorted this differentiation trajectory (Fig 6A, right panel). The myeloid and stem-like arms were disorganised, with cells from different clusters scattered over the trajectory, although the erythroid branch appeared unaffected. Our results are consistent with RUNX1-ETO halting differentiation with cells forming a continuum of mixed differentiation stages rather than a trajectory, which would be in agreement with the fact that cell cycle is blocked and lineage fate decisions cannot be properly executed. The same scenario can be seen when the expression of specific genes in the different cell populations is projected on the trajectory (Fig 6B). The expression of genes such as *GATA2* and *SPI1 (PU.1)* scattered all over the trajectory after induction (Fig 6B) and their expression was down-regulated in stem-like (orange) and 5-Dox enriched (green) populations (Fig 6B, Fig S5). However, the analysis of *SPI1* (PU.1) expression in monocytic cells (purple) showed a different picture (Fig 6B, S5), as *SPI1* appeared to be much less perturbed by RUNX1-ETO induction, suggesting that cells that have passed a certain differentiation state become less sensitive to perturbation. This might be explained due to additional enhancers that are not dependent on prior expression of RUNX1 (Leddin et al., 2011). In contrast, *SOX4* upregulation is confined to the stem-like and the 5-Dox induced cell population (Fig S5).

**FIGURE 6.**
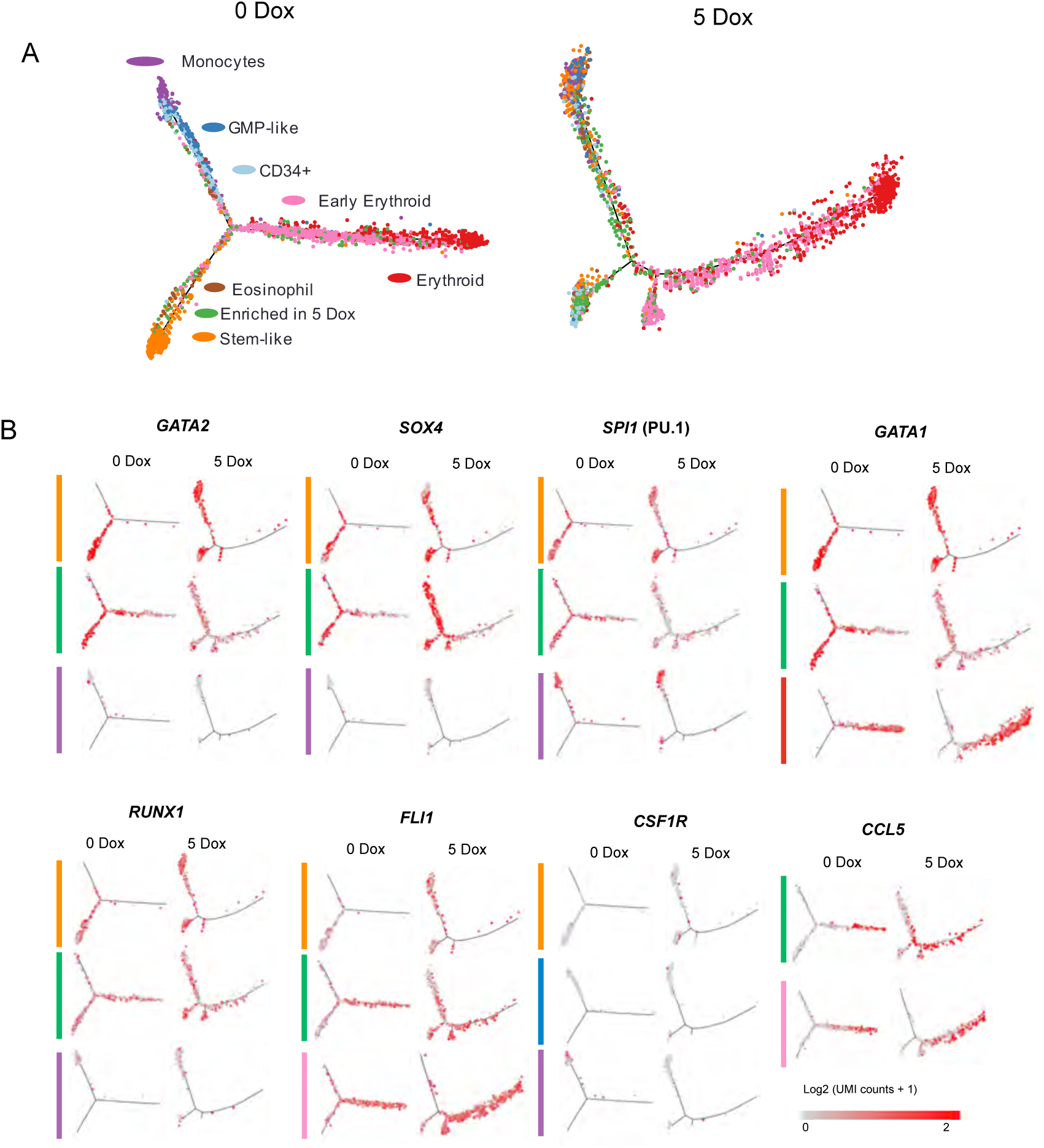
RUNX1-ETO induction distorts the myeloid but not the erythroid differentiation trajectory and dysregulates genes involved in stem/progenitor development. A) Trajectory analysis using the Monocle algorithm of the sorted cell populations plotted according to each cell cluster. B) Expression of individual marker genes projected on the trajectories plotted according to each cell cluster in (A). Colour intensity represents number of transcripts sequenced in Log2 of unique molecular identifier (UMI) counts +1.

Taken together, our results are consistent with the finding that RUNX1-ETO reprograms a specific, strongly responsive early myeloid cell population, leading to the dysregulation of genes involved in stem/progenitor cell development and arrested differentiation as well as growth of these cells.

## Discussion

### RUNX1-ETO induction at balanced levels in human progenitor cells induces quiescence and promotes their survival

The t(8;21) translocation can be observed *in utero* and *RUNX1-ETO*-expressing clones can be detected in post-natal blood samples, suggesting that cells that acquired the mutation might form a pre-leukemic clonal reservoir (Wiemels et al., 2010). Our laboratories have established a protocol for the generation of CD34+ RUNX1C+ definitive hematopoietic progenitors arising from *HOXA+* hemogenic vasculature, that resemble cells arising during human intra-embryonic hematopoiesis within the AGM region (Ng et al., 2016). These progenitors present similar transcriptional profiles, cell surface receptors and signaling molecules to cells sorted from human AGM. The work presented here demonstrates that such cells also display an accessible chromatin landscape resembling the HSC/MPP pattern found in adult hematopoietic cells.

In agreement with previous experiments in murine ESCs (Regha et al., 2015), *RUNX1-ETO* expression at a high levels in differentiating human ESCs abrogated blood formation and here we show that it also perturbs vasculogenesis, similar to what has been shown in transgenic mice (Yergeau et al., 1997). Expression of *RUNX1-ETO* before the EHT, even at an equivalent level to that of endogenous *RUNX1*, caused a dramatic disorganization of vascular structures and formation of morphologically abnormal hematopoietic progenitors. However, induction of the same level of RUNX1-ETO after the EHT allowed the formation of progenitor cells, but promoted the accumulation of a cell population expressing markers of immature blood progenitors with a CD34+CD38-CD90+ phenotype. This CD34+CD38-CD90+CD45+RUNX1+ signature in our *RUNX1-ETO*-expressing progenitors is shared with an embryonic population of cells containing the first few definitive human HSCs (Ivanovs et al., 2014).

Balanced levels of RUNX1-ETO expression (i) do not block blood formation, (ii) confer survival but not proliferation to a subset of progenitors, (iii) do not cause apoptosis (data not shown) and (vi) maintain cells in a quiescent stage. Moreover, the latent colony forming activity of progenitors that had been previously rendered quiescent by RUNX1-ETO induction re-emerged upon removal of Dox from the methylcellulose CFU assay. Indeed, these cells exhibited a higher yield of colonies when compared to their uninduced counterparts, suggesting that the blocked differentiation state is reversible even after extended *RUNX1-ETO* expression in culture. This result is consistent with prior re-plating assays of RUNX-ETO-expressing mouse bone marrow cells which showed an increase of self-renewal but not proliferation (Rhoades et al., 2000). Our observations are also consistent with the idea of the existence of a reservoir of cells harbouring the t(8;21) translocation in a quiescent pre-leukemic state, as reported by the presence of a small population of HSC harbouring the translocation in t(8;21) patients in remission (Shima et al., 2014). In contrast, studies using retroviral transduction of human CD34+ cord blood hematopoietic cells to constitutively express *RUNX1-ETO* were reported to enhance *in vitro* proliferation, whilst maintaining self-renewal and differentiation capacity (Mulloy et al., 2002, 2003). These discrepancies might be explained by differences in the expression levels of *RUNX1-ETO* used and/or by selection of specific clones of cells that were outgrowing in culture (Mulloy et al., 2002, 2003), indicating that the growth arrest can be surpassed. A dual role of RUNX1-ETO in blocking differentiation and arresting cell growth has been previously described (Burel et al., 2001), although they observed that RUNX1-ETO-dependent growth arrest resulted in apoptosis. It is important to note that those studies were conducted in a cell line (U937) representing a different type of AML, and probably carrying additional oncogenic mutations, suggesting that RUNX1-ETO cannot reprogram one leukemic cell type into another. This observation is consistent with previous work from our lab that showed that each leukemogenic mutation directs hematopoietic precursors through a different developmental path (Assi et al., 2019).

### RUNX1-ETO induction causes extensive global chromatin reorganisation and shuts down the RUNX1-directed gene expression program

We found that the pattern of up and down-regulated RUNX1-ETO target genes is tightly correlated to changes observed in t(8;21) patient cells. Differential gene expression analyses upon induction of RUNX1-ETO showed downregulation of myelopoiesis, cell cycle and DNA repair genes. This finding is consistent with the expression of surface markers characteristic of immature cells in RUNX1-ETO-induced progenitors and the lack of proliferation of such cells. Interestingly, RUNX1-ETO induction caused an upregulation of genes from multiple signalling pathways, such as the MAPK pathway. General signaling pathway activation could occur as a survival response from the cells trying to compensate for the RUNX1-ETO-mediated block on the cell cycle. This hypothesis is supported by data from established t(8;21) AML cells that show that high *CDK6* and *CCND1/2* expression levels are dependent on the AP-1 transcription factor family, which mediates MAPK signalling and whose expression, in turn, depends on the presence of RUNX1-ETO (Martinez-Soria et al., 2019). Moreover, AP-1 is a critical factor for the establishment of t(8;21) AML in xenotransplantation experiments (Assi et al., 2019). In order to develop overt disease, RUNX1-ETO cooperates with mutated growth factor receptors, such as KIT or FLT3 (Schessl et al., 2005; Wichmann et al., 2015), indicating that such mutations may act upon an already partially activated signalling landscape. So far, we have not been able to grow RUNX1-ETO-transduced human cells without an extensive selection and outgrowth in culture (unpublished results), indicating that we do not yet understand which signals are essential and responsible for the epigenetic drift that permits growth *in vitro*.

Changes in gene expression after induction of RUNX1-ETO were mediated by rapid reorganisation of the epigenetic landscape, leading to loss of accessibility of thousands of sites enriched in RUNX motifs and bound by RUNX1, although *RUNX1* expression per se was not affected. Concomitantly, H3K27 acetylation was reduced, demonstrating that binding of RUNX1-ETO directly interferes with that of RUNX1. Interestingly, these changes were most pronounced at distal elements. It is now clear that cellular identity is encoded within the combination of active distal elements (Corces et al., 2016; Heinz et al., 2010), which demonstrates that RUNX1-ETO profoundly and rapidly interferes with the identity of its target cells.

It has been shown that t(8;21) cells are absolutely dependent on the presence of a normal *RUNX1* allele, which is required to balance the detrimental effects of RUNX1-ETO expression by upregulating critical mitotic checkpoint genes (Ben-Ami et al., 2013; Loke et al., 2017). It is therefore possible that restoring the growth of RUNX1-ETO-induced cells requires the re-establishment of, at least, part of the RUNX1-mediated gene expression program. This is likely to include the expression of *SPI1* (PU.1) (Huang et al., 2008) or *CEBPA*, but also genes that override the RUNX1-ETO mediated cell cycle block.

### RUNX1-ETO expression blocks the differentiation of a specific cell population

Reprogramming of one cell fate into another requires a complete reorganisation of the epigenome and is facilitated by reprogramming cells within a similar developmental pathway (Graf and Enver, 2009), but can also be enforced upon cells from different pathways by overexpression of complementary transcription factors (Takahashi and Yamanaka, 2006). One of the most important and unanswered questions in AML research is the understanding of the nature of the target cells that are susceptible to transformation in order to understand the first response to an oncogenic hit. This notion is exemplified by our finding that RUNX1C- and RUNX1C+ cells significantly differ in their response to the expression of the fusion protein. Our scRNA-Seq data clearly demonstrate that only early multipotent myeloid precursors respond to RUNX1-ETO induction. Moreover, those are the only cells suffering a down-regulation of the myeloid master regulator PU.1, without which myelopoiesis is strongly perturbed (McKercher et al., 1996; Scott et al., 1994), providing a molecular explanation for the RUNX1-ETO-mediated differentiation block. Expression of *RUNX1-ETO* also leads to an upregulation of *SOX4* in the 5-Dox enriched cell cluster, which is consistent with the down-regulation of *CEBPA. SOX4* expression is required for self-renewal of HSCs as well as LSCs, and its expression has been shown to be upregulated in HSCs from *CEBPA*-null mice and in patients with abnormal C/EBPα function (Zhang et al., 2013), and thus with myelopoiesis.

In summary, we have shown that we can use the differentiation of human ES cells into definitive hematopoietic progenitor cells to gain insight into the earliest events of epigenetic reprogramming by RUNX1-ETO and its interplay with RUNX1 in a human embryonic setting. Our future experiments are aimed at further understanding the nature of the growth stimulus required to overcome RUNX1-ETO-mediated growth arrest.

## Experimental Procedures

### Generation and validation of targeted inducible RUNX1-ETO SOX17^mCHERRY/w^ RUNX1C^GFP/w^ hESC lines

The dual reporter SOX17^mCHERRY/w^RUNX1C^GFP/w^ hESC H9 line was previously generated by us (Ng et al., 2016). RUNX1-ETO-AAVS1 targeting vectors comprised a 804-bp 5’ homology arm, a tetracycline-inducible promoter (TRE3G-CMV) driving expression of HA-RUNX1-ETO cDNA sequence, Puromycin N-acetyltransferase resistance cassette with gene expression to be driven by endogenous promoter after genomic insertion, a CAG promoter driving expression of a modified Tet3G reverse tetracycline-controlled transactivator (rtTA) and a 837-bp 3’ homology arm. Vectors were electroporated with a pair of AAVS1 Transcription Activator-Like Effector Nucleases (TALENs) into SOX17^mCHERRY/w^RUNX1C^GFP/w^ hESC H9 cells, which were then selected for Puromycin-resistant colony growth. Single cell sorted clones were screened for transgene insertion by PCR using primers designed to amplify the boundaries of the genomic insertion. For other details see Supplementary methods.

### Maintenance and hematopoietic differentiation of hESC

Culture and enzymatic passaging of hESC lines was conducted as previously reported (Ng et al., 2008). Hematopoietic differentiation of hPSCs was performed following the spin Embryo Body (EB) method in STAPEL medium (Ng et al., 2008) with supplements used as reported (Ng et al., 2016) with some modifications, detailed in Supplemental methods.

### Colony-forming assays

Colony-forming-unit (CFU) assays were performed as reported (Ng et al., 2016) with some modifications. Briefly, 3-5 x 10^3^ cells were cultured in 1% methylcellulose, supplemented with the ‘5-factor’ cytokine mix (as used in the hematopoietic differentiation) plus 10 μg/ml human low-density lipoproteins (hLDL, Stem Cell Technologies cat# 02698) and 5 U/ml erythropoietin (EPO, PeproTech cat# 100-64). For the preparation of 1% methylcellulose, 40 ml serum-free 2.6% MethoCult^TM^ H4100 (Stem Cell Technologies #01400) was mixed with an equal volume of 2x STAPEL-P medium (STAPEL medium made with IMDM containing 2x supplements and without PFHMII) plus 20 ml of 1x STAPEL medium to give a final volume of 100 ml. Cells were cultured either with or without Dox and each condition was set up in triplicates in ultra-low attachment 24-well plates (cat# NUN144530). Plates were scored for hematopoietic CFUs after 7 to 10 days.

### Replating assays

Replating assays were conducted on non-adherent hematopoietic progenitors plated at a known concentration on Matrigel-coated wells of a 6-well plate. Cells were harvested, counted and replated weekly. Live and death cells were determined using a FL Countess-II Automated Cell Counter (Thermo Fisher Scientific) after Tripan-Blue staining.

### RNA-Seq library preparation

RNA-sequencing (seq) libraries were prepared using a TruSeq® Stranded mRNA Library Prep (Illumina cat# 20020594) following the Low Sample (LS) workflow according to manufacturer’s instructions. Libraries were subjected to a quality control using a High Sensitivity DNA chip on an Agilent Technologies 2100 Bioanalyser™ instrument and were quantified using the RT-qPCR-based method KAPA Library Quantification Kit for Illumina Sequencing Platforms (Roche cat# KR0405), following the manufacturer’s protocol. Libraries were run in a pool of twelve indexed libraries in a NextSeq (Illumina) machine using sequencing by synthesis chemistry and a NextSeq® 500/550 High Output 150 cycle sequencing kit v2 (cat# FC-404-2002), obtaining 75 bp paired-end reads.

### Assay for transposase-accessible chromatin with high-throughput sequencing (ATAC-seq)

Chromatin accessibility was evaluated by ATAC-seq using a modified protocol to as reported (Buenrostro et al., 2015; Corces et al., 2016), Briefly, 50,000 single-cell sorted hematopoietic progenitors were pelleted and snap-frozen upon resuspension in 5 μl sucrose freezing buffer, consisting of 60 mM KCl, 15 mM NaCl, 5 mM MgCl_2_, 10 mM Tris pH 7.4 and 1.5 M sucrose. Transposition reaction was performed for 30 minutes at 37°C upon addition of 45 μl of ATAC reaction mix consisting of 25 μl Tagmentation DNA Buffer (Illumina cat# FC-121-1030, Nextera DNA Library Prep Kit), 2.5 μl Tn5 Transposase enzyme (Illumina cat# FC-121-1030, Nextera DNA Library Prep Kit), 1 μl of 0.5% Digitonin (Promega cat# G9441) and 16.5 μl water. For other details see Supplemental methods.

### Chromatin Immunoprecipitation with high-throughput sequencing (ChIP-Seq)

ChIP was performed on CD34+ magnetic sorted samples (RUNX1-ETO and RUNX) and on non-adhered floating mixed progenitors (H3K4me3 and H3K27ac) after single crosslink, as reported (Obier et al., 2016).

### Single Cell RNA-Seq (scRNA-Seq)

Non-adherent progenitors at day 22 of differentiation (untreated and 24-hour Dox treatment) were sorted for CD45+ CD34+ and RUNX1C+. Cells were re-suspended in 80 μl at a concentration of 100-1200 cells/μl for evaluation of cell viability prior to loading of 4000 single cells on a Chromium Single Cell Instrument (10X Genomics). Library generation for scRNA-seq was performed by the Genomics Birmingham Sequencing Facility using the Chromium Single Cell 3′ Library and Gel Bead Kit v2 (10X Genomics). Libraries were paired-end sequenced on an Illumina NextSeq machine using the cycle parameters recommended by 10X Genomics.

### Statistical analysis

Experiments were analysed using GraphPad Prism versions 5–7 (GraphPad Software Inc.) and Microsoft Excel (Microsoft corporation).

### Bioinformatic Data processing and analysis

Details of data analysis iare described in Supplemental Methods.

## Author Contributions

M.N. performed experiments and wrote the paper, P.K. analysed the data, E.S.N. and E.G.S. helped with experiments, A.G.E. and C.B. conceived the study and wrote the paper.

## Acknowledgments

This work was funded by a U21 Melbourne studentship grant to the University of Birmingham held by Monica Nafria, and a grant from the Birmingham CRUK Centre to C.B. We thank the Birmingham Next Generation Sequencing Facility (Genomics Birmingham) for expert service. Monica Nafria was awarded The Henry and Rachael Ackman Travelling Scholarship by the University of Melbourne Faculty of Medicine, Dentistry and Health Sciences. Work in the laboratories of A.G.E and E.G.S was funded by the National Health and Medical Research Council of Australia (GNT1068866, GNT1129861, GNT1138717), by the Australian Research Council Special Research Initiative in Stem Cells (Stem Cells Australia), the Children’s Cancer Foundation, and by the Stafford Fox Medical Research Foundation. E.G.S. (GNT1079004) and A.G.E. (GNT1117596) are Research Fellows of the National Health and Medical Research Council (Australia). Additional infrastructure funding to the Murdoch Children’s Research Institute was provided by the Australian Government National Health and Medical Research Council Independent Research Institute Infrastructure Support Scheme and the Victorian Government’s Operational Infrastructure Support Program.

## References

Assi, S.A., Imperato, M.R., Coleman, D.J.L., Pickin, A., Potluri, S., Ptasinska, A., Chin, P.S., Blair, H., Cauchy, P., James, S.R., et al. (2019). Subtype-specific regulatory network rewiring in acute myeloid leukemia. Nat. Genet. 51, 151–162.

Ben-Ami, O., Friedman, D., Leshkowitz, D., Goldenberg, D., Orlovsky, K., Pencovich, N., Lotem, J., Tanay, A., and Groner, Y. (2013). Addiction of t(8;21) and inv(16) Acute Myeloid Leukemia to Native RUNX1. Cell Rep. 4, 1131–1143.

Bertrand, J.Y., Chi, N.C., Santoso, B., Teng, S., Stainier, D.Y.R., and Traver, D. (2010). Haematopoietic stem cells derive directly from aortic endothelium during development. Nature 464, 108–111.

Bindea, G., Mlecnik, B., Hackl, H., Charoentong, P., Tosolini, M., Kirilovsky, A., Fridman, W.-H., Pagès, F., Trajanoski, Z., and Galon, J. (2009). ClueGO: a Cytoscape plug-in to decipher functionally grouped gene ontology and pathway annotation networks. Bioinformatics 25, 1091–1093.

Boisset, J.-C., van Cappellen, W., Andrieu-Soler, C., Galjart, N., Dzierzak, E., and Robin, C. (2010). In vivo imaging of haematopoietic cells emerging from the mouse aortic endothelium. Nature 464, 116–120.

Bolger, A.M., Lohse, M., and Usadel, B. (2014). Trimmomatic: a flexible trimmer for Illumina sequence data. Bioinformatics 30, 2114–2120.

Bomken, S., Buechler, L., Rehe, K., Ponthan, F., Elder, A., Blair, H., Bacon, C.M., Vormoor, J., and Heidenreich, O. (2013). Lentiviral marking of patient-derived acute lymphoblastic leukaemic cells allows in vivo tracking of disease progression. Leukemia 27, 1792.

Bonifer, C., and Cockerill, P.N. (2017). Chromatin priming of genes in development: Concepts, mechanisms and consequences. Exp. Hematol. 49, 1–8.

Boyd, A.L., Aslostovar, L., Reid, J., Ye, W., Tanasijevic, B., Porras, D.P., Shapovalova, Z., Almakadi, M., Foley, R., Leber, B., et al. (2018). Identification of Chemotherapy-Induced Leukemic-Regenerating Cells Reveals a Transient Vulnerability of Human AML Recurrence. Cancer Cell 34, 483–498.e5.

De Bruijn, M.F.T.R., Ma, X., Robin, C., Ottersbach, K., Sanchez, M., and Dzierzak, E. (2002). Haematopoietic Stem Cells Localize to the Endothelial Cell Layer in the Midgestation Mouse Aorta. Immunity 16, 673–683.

Buenrostro, J.D., Wu, B., Chang, H.Y., and Greenleaf, W.J. (2015). ATAC-seq: A method for assaying chromatin accessibility genome-wide. Curr. Protoc. Mol. Biol. 2015, 21.29.1–21.29.9.

Burel, S.A., Harakawa, N., Zhou, L., Pabst, T., Tenen, D.G., and Zhang, D.E. (2001). Dichotomy of AML1-ETO functions: growth arrest versus block of differentiation. Mol. Cell. Biol. 21, 5577–5590.

Butler, A., Hoffman, P., Smibert, P., Papalexi, E., and Satija, R. (2018). Integrating single-cell transcriptomic data across different conditions, technologies, and species. Nat. Biotechnol. 36, 411–420.

Cabezas-Wallscheid, N., Eichwald, V., de Graaf, J., Löwer, M., Lehr, H.-A., Kreft, A., Eshkind, L., Hildebrandt, A., Abassi, Y., Heck, R., et al. (2013). Instruction of haematopoietic lineage choices, evolution of transcriptional landscapes and cancer stem cell hierarchies derived from an AML1-ETO mouse model. EMBO Mol. Med. 5, 1804–1820.

Clarke, R.L., Yzaguirre, A.D., Yashiro-Ohtani, Y., Bondue, A., Blanpain, C., Pear, W.S., Speck, N. a, and Keller, G. (2013). The expression of Sox17 identifies and regulates haemogenic endothelium. Nat. Cell Biol. 15, 502–510.

Corces, M.R., Buenrostro, J.D., Wu, B., Greenside, P.G., Chan, S.M., Koenig, J.L., Snyder, M.P., Pritchard, J.K., Kundaje, A., Greenleaf, W.J., et al. (2016). Lineage-specific and single-cell chromatin accessibility charts human hematopoiesis and leukemia evolution. Nat. Genet. 48, 1193–1203.

Graf, T., and Enver, T. (2009). Forcing cells to change lineages. Nature 462, 587–594.

Heinz, S., Benner, C., Spann, N., Bertolino, E., Lin, Y.C., Laslo, P., Cheng, J.X., Murre, C., Singh, H., and Glass, C.K. (2010). Simple Combinations of Lineage-Determining Transcription Factors Prime cis-Regulatory Elements Required for Macrophage and B Cell Identities. Mol. Cell 38, 576–589.

Higuchi, M., O’Brien, D., Kumaravelu, P., Lenny, N., Yeoh, E.-J., and Downing, J.R. (2002). Expression of a conditional AML1-ETO oncogene bypasses embryonic lethality and establishes a murine model of human t(8;21) acute myeloid leukemia. Cancer Cell 1, 63–74.

Huang, G., Zhang, P., Hirai, H., Elf, S., Yan, X., Chen, Z., Koschmieder, S., Okuno, Y., Dayaram, T., Growney, J.D., et al. (2008). PU.1 is a major downstream target of AML1 (RUNX1) in adult mouse hematopoiesis. Nat. Genet. 40, 51–60.

Ivanovs, A., Rybtsov, S., Anderson, R.A., Turner, M.L., and Medvinsky, A. (2014). Identification of the Niche and Phenotype of the First Human Hematopoietic Stem Cells. Stem Cell Reports 2, 449–456.

Jaffredo, T., Gautier, R., Eichmann, A., and Dieterlen-Lièvre, F. (1998). Intraaortic Hemopoietic Cells are Derived from Endothelial Cells During Ontogeny. Development 125, 4575–4583.

Karolchik, D., Hinrichs, A.S., Furey, T.S., Roskin, K.M., Sugnet, C.W., Haussler, D., and Kent, W.J. (2004). The UCSC Table Browser data retrieval tool. Nucleic Acids Res. 32, 493D–496.

Kim, D., Langmead, B., and Salzberg, S.L. (2015). HISAT: a fast spliced aligner with low memory requirements. Nat. Methods 12, 357–360.

Kissa, K., and Herbomel, P. (2010). Blood stem cells emerge from aortic endothelium by a novel type of cell transition. Nature 464, 112–115.

Lam, E.Y.N., Hall, C.J., Crosier, P.S., Crosier, K.E., and Flores, M.V. (2010). Live imaging of Runx1 expression in the dorsal aorta tracks the emergence of blood progenitors from endothelial cells. Blood 116, 909–914.

Langfelder, P., Zhang, B., and Horvath, S. (2008). Defining clusters from a hierarchical cluster tree: the Dynamic Tree Cut package for R. Bioinformatics 24, 719–720.

Langmead, B., and Salzberg, S.L. (2012). Fast gapped-read alignment with Bowtie 2. Nat. Methods 9, 357–359.

Leddin, M., Perrod, C., Hoogenkamp, M., Ghani, S., Assi, S., Heinz, S., Wilson, N.K., Follows, G., Schönheit, J., Vockentanz, L., et al. (2011). Two distinct auto-regulatory loops operate at the PU.1 locus in B cells and myeloid cells. Blood 117, 2827–2838.

Loke, J., Assi, S.A., Imperato, M.R., Ptasinska, A., Cauchy, P., Grabovska, Y., Soria, N.M., Raghavan, M., Delwel, H.R., Cockerill, P.N., et al. (2017). RUNX1-ETO and RUNX1-EVI1 Differentially Reprogram the Chromatin Landscape in t(8;21) and t(3;21) AML. Cell Rep. 19, 1654–1668.

Lutterbach, B., Westendorf, J.J., Linggi, B., Patten, A., Moniwa, M., Davie, J.R., Huynh, K.D., Bardwell, V.J., Lavinsky, R.M., Rosenfeld, M.G., et al. (1998). ETO, a target of t(8;21) in acute leukemia, interacts with the N-CoR and mSin3 corepressors. Mol. Cell. Biol. 18, 7176–7184.

Mandoli, A., Singh, A.A., Prange, K.H.M., Tijchon, E., Oerlemans, M., Dirks, R., Ter Huurne, M., Wierenga, A.T.J., Janssen-Megens, E.M., Berentsen, K., et al. (2016). The Hematopoietic Transcription Factors RUNX1 and ERG Prevent AML1-ETO Oncogene Overexpression and Onset of the Apoptosis Program in t(8;21) AMLs. Cell Rep. 17, 2087–2100.

Martens, J.H. a., Mandoli, a., Simmer, F., Wierenga, B.-J., Saeed, S., Singh, a. a., Altucci, L., Vellenga, E., and Stunnenberg, H.G. (2012). ERG and FLI1 binding sites demarcate targets for aberrant epigenetic regulation by AML1-ETO in acute myeloid leukemia. Blood 120, 4038–4048.

Martinez-Soria, N., McKenzie, L., Draper, J., Ptasinska, A., Issa, H., Potluri, S., Blair, H.J., Pickin, A., Isa, A., Chin, P.S., et al. (2019). The Oncogenic Transcription Factor RUNX1/ETO Corrupts Cell Cycle Regulation to Drive Leukemic Transformation. Cancer Cell 35, 705.

McKercher, S.R., Torbett, B.E., Anderson, K.L., Henkel, G.W., Vestal, D.J., Baribault, H., Klemsz, M., Feeney, a J., Wu, G.E., Paige, C.J., et al. (1996). Targeted disruption of the PU.1 gene results in multiple hematopoietic abnormalities. EMBO J. 15, 5647–5658.

Miyamoto, T., Weissman, I.L., and Akashi, K. (2000). AML1/ETO-expressing nonleukemic stem cells in acute myelogenous leukemia with 8;21 chromosomal translocation. Proc. Natl. Acad. Sci. U. S. A. 97, 7521–7526.

Miyoshi, H., Shimizu, K., Kozu, T., Maseki, N., Kaneko, Y., and Ohki, M. (1991). t(8;21) breakpoints on chromosome 21 in acute myeloid leukemia are clustered within a limited region of a single gene, AML1. Proc. Natl. Acad. Sci. 88, 10431–10434.

Mulloy, J.C., Mackenzie, K.L., Berguido, F.J., Moore, M. a S., and Nimer, S.D. (2002). The AML1-ETO fusion protein promotes the expansion of human hematopoietic stem cells. Blood 99, 15–23.

Mulloy, J.C., Cammenga, J., Berguido, F.J., Wu, K., Zhou, P., Comenzo, R.L., Jhanwar, S., Moore, M. a. S., and Nimer, S.D. (2003). Maintaining the self-renewal and differentiation potential of human CD34+ hematopoietic cells using a single genetic element. Blood 102, 4369.

Ng, E.S., Davis, R., Stanley, E.G., and Elefanty, A.G. (2008). A protocol describing the use of a recombinant protein-based, animal product-free medium (APEL) for human embryonic stem cell differentiation as spin embryoid bodies. Nat. Protoc. 3, 768–776.

Ng, E.S., Azzola, L., Bruveris, F.F., Calvanese, V., Phipson, B., Vlahos, K., Hirst, C., Jokubaitis, V.J., Yu, Q.C., Maksimovic, J., et al. (2016). Differentiation of human embryonic stem cells to HOXA+ hemogenic vasculature that resembles the aorta-gonad-mesonephros. Nat. Biotechnol. 34, 1168–1179.

North, T.E., De Bruijn, M.F.T.R., Stacy, T., Talebian, L., Lind, E., Robin, C., Binder, M., Dzierzak, E., and Speck, N. a. (2002). Runx1 expression marks long-term repopulating hematopoietic stem cells in the midgestation mouse embryo. Immunity 16, 661–672.

O’Leary, N.A., Wright, M.W., Brister, J.R., Ciufo, S., Haddad, D., McVeigh, R., Rajput, B., Robbertse, B., Smith-White, B., Ako-Adjei, D., et al. (2016). Reference sequence (RefSeq) database at NCBI: current status, taxonomic expansion, and functional annotation. Nucleic Acids Res. 44, D733–D745.

Obier, N., Cauchy, P., Assi, S.A., Gilmour, J., Lie-A-Ling, M., Lichtinger, M., Hoogenkamp, M., Noailles, L., Cockerill, P.N., Lacaud, G., et al. (2016). Cooperative binding of AP-1 and TEAD4 modulates the balance between vascular smooth muscle and hemogenic cell fate. Development 143, 4324–4340.

Pertea, M., Pertea, G.M., Antonescu, C.M., Chang, T.-C., Mendell, J.T., and Salzberg, S.L. (2015). StringTie enables improved reconstruction of a transcriptome from RNA-seq reads. Nat. Biotechnol. 33, 290–295.

Ptasinska, A., Assi, S.A., Mannari, D., James, S.R., Williamson, D., Dunne, J., Hoogenkamp, M., Wu, M., Care, M., McNeill, H., et al. (2012). Depletion of RUNX1/ETO in t(8;21) AML cells leads to genome-wide changes in chromatin structure and transcription factor binding. Leukemia 26, 1829–1841.

Ptasinska, A., Assi, S.A., Martinez-Soria, N., Imperato, M.R., Piper, J., Cauchy, P., Pickin, A., James, S.R., Hoogenkamp, M., Williamson, D., et al. (2014). Identification of a dynamic core transcriptional network in t(8;21) AML that regulates differentiation block and self-renewal. Cell Rep. 8, 1974–1988.

Qian, K., Huang, C.T.-L., Huang, C.-L., Chen, H., Blackbourn, L.W., Chen, Y., Cao, J., Yao, L., Sauvey, C., Du, Z., et al. (2014). A simple and efficient system for regulating gene expression in human pluripotent stem cells and derivatives. Stem Cells 32, 1230–1238.

Qiu, X., Mao, Q., Tang, Y., Wang, L., Chawla, R., Pliner, H.A., and Trapnell, C. (2017). Reversed graph embedding resolves complex single-cell trajectories. Nat. Methods 14, 979–982.

Ramírez, F., Ryan, D.P., Grüning, B., Bhardwaj, V., Kilpert, F., Richter, A.S., Heyne, S., Dündar, F., and Manke, T. (2016). deepTools2: a next generation web server for deep-sequencing data analysis. Nucleic Acids Res. 44, W160–W165.

Regha, K., Assi, S. a., Tsoulaki, O., Gilmour, J., Lacaud, G., and Bonifer, C. (2015). Developmental-stage-dependent transcriptional response to leukaemic oncogene expression. Nat. Commun. 6, 7203.

Rhoades, K.L., Hetherington, C.J., Harakawa, N., Yergeau, D. a, Zhou, L., Liu, L.Q., Little, M.T., Tenen, D.G., and Zhang, D.E. (2000). Analysis of the role of AML1-ETO in leukemogenesis, using an inducible transgenic mouse model. Blood 96, 2108–2115.

Ritchie, M.E., Phipson, B., Wu, D., Hu, Y., Law, C.W., Shi, W., and Smyth, G.K. (2015). limma powers differential expression analyses for RNA-sequencing and microarray studies. Nucleic Acids Res. 43, e47–e47.

Saldanha, A.J. (2004). Java Treeview--extensible visualization of microarray data. Bioinformatics 20, 3246–3248.

Schessl, C., Rawat, V.P.S., Cusan, M., Deshpande, A., Kohl, T.M., Rosten, P.M., Spiekermann, K., Humphries, R.K., Schnittger, S., Kern, W., et al. (2005). The AML1-ETO fusion gene and the FLT3 length mutation collaborate in inducing acute leukemia in mice. J. Clin. Invest. 115, 2159–2168.

Scott, E., Simon, M., Anastasi, J., and Singh, H. (1994). Requirement of transcription factor PU.1 in the development of multiple hematopoietic lineages. Science (80-.). 265, 1573–1577.

Shannon, P., Markiel, A., Ozier, O., Baliga, N.S., Wang, J.T., Ramage, D., Amin, N., Schwikowski, B., and Ideker, T. (2003). Cytoscape: a software environment for integrated models of biomolecular interaction networks. Genome Res. 13, 2498–2504.

Shima, T., Miyamoto, T., Kikushige, Y., Yuda, J., Tochigi, T., Yoshimoto, G., Kato, K., Takenaka, K., Iwasaki, H., Mizuno, S., et al. (2014). The ordered acquisition of Class II and Class I mutations directs formation of human t (8; 21) acute myelogenous leukemia stem cell. Exp. Hematol. 42, 955–965.e5.

Sroczynska, P., Lancrin, C., Kouskoff, V., and Lacaud, G. (2009). The differential activities of Runx1 promoters define milestones during embryonic hematopoiesis. Blood 114, 5279–5289.

Subramanian, A., Tamayo, P., Mootha, V.K., Mukherjee, S., Ebert, B.L., Gillette, M.A., Paulovich, A., Pomeroy, S.L., Golub, T.R., Lander, E.S., et al. (2005). Gene set enrichment analysis: a knowledge-based approach for interpreting genome-wide expression profiles. Proc. Natl. Acad. Sci. U. S. A. 102, 15545–15550.

Takahashi, K., and Yamanaka, S. (2006). Induction of Pluripotent Stem Cells from Mouse Embryonic and Adult Fibroblast Cultures by Defined Factors. Cell 126, 663–676.

Trapnell, C., Cacchiarelli, D., Grimsby, J., Pokharel, P., Li, S., Morse, M., Lennon, N.J., Livak, K.J., Mikkelsen, T.S., and Rinn, J.L. (2014). The dynamics and regulators of cell fate decisions are revealed by pseudotemporal ordering of single cells. Nat. Biotechnol. 32, 381–386.

Wichmann, C., Quagliano-Lo Coco, I., Yildiz, Ö., Chen-Wichmann, L., Weber, H., Syzonenko, T., Döring, C., Brendel, C., Ponnusamy, K., Kinner, A., et al. (2015). Activating c-KIT mutations confer oncogenic cooperativity and rescue RUNX1/ETO-induced DNA damage and apoptosis in human primary CD34+ hematopoietic progenitors. Leukemia 29, 279–289.

Wiemels, J.L., Xiao, Z., Buffler, P. a, Maia, A.T., Ma, X., Dicks, B.M., Martyn, T., Zhang, L., Feusner, J., Wiencke, J., et al. (2010). In utero origin of t(8;21) AML1-ETO translocations in childhood acute myeloid leukemia. Neoplasia 99, 3801–3805.

Yergeau, D.A., Hetherington, C.J., Wang, Q., Zhang, P., Sharpe, A.H., Binder, M., Marín-Padilla, M., Tenen, D.G., Speck, N.A., and Zhang, D.E. (1997). Embryonic lethality and impairment of haematopoiesis in mice heterozygous for an AML1-ETO fusion gene. Nat. Genet. 15, 303–306.

Yuan, Y., Zhou, L., Miyamoto, T., Iwasaki, H., Harakawa, N., Hetherington, C.J., Burel, S. a, Lagasse, E., Weissman, I.L., Akashi, K., et al. (2001). AML1-ETO expression is directly involved in the development of acute myeloid leukemia in the presence of additional mutations. Proc. Natl. Acad. Sci. U. S. A. 98, 10398–10403.

Zhang, H., Alberich-Jorda, M., Amabile, G., Yang, H., Staber, P.B., Di Ruscio, A., Diruscio, A., Welner, R.S., Ebralidze, A., Zhang, J., et al. (2013). Sox4 is a key oncogenic target in C/EBPα mutant acute myeloid leukemia. Cancer Cell 24, 575–588.

Zhang, Y., Liu, T., Meyer, C.A., Eeckhoute, J., Johnson, D.S., Bernstein, B.E., Nussbaum, C., Myers, R.M., Brown, M., Li, W., et al. (2008). Model-based Analysis of ChIP-Seq (MACS). Genome Biol. 9, R137.

