## Supplemental figures, figure legends, materials and methods for "RUNX1-ETO induction rapidly alters chromatin landscape and growth of a specific sub-population of hESC-derived myeloid precursor cells by interfering with RUNX1 regulation"

#### Supplemental Materials

##### List of Supplemental files

1. Supplemental Figures and Figure legends

2. Supplemental methods

3. Supplemental Tables

4. Supplemental data-sets

Supplemental Dataset 1: Alignment statistics

Supplemental Dataset 2: Gene expression levels and differentia gene expression in RUNX1C+ and RUNX1C- cells. Related to Figure 3A

Supplemental Dataset 3: Dose response of groups of genes in RUNX1C+ and RUNX1C- cells. Related to Figure S2E

Supplemental Dataset 4: KEGG pathways of genes altered by RUNX1-ETO induction in RUNX1C+ and RUNX1C- cells. Related to Figure 3B.

Supplemental Dataset 5: List of up- and down-regulated RUNX1-ETO and RUNX1 target genes. Related to Figure S3E,F

Supplemental Dataset 6: List of marker genes used to classify single cell populaitons. Related to Figure S4B

Supplemental Dataset 7: List of genes associated with KEGG pathways shown in Figure S6B.

### Supplemental Figures and Figure Legends

#### Figure S1

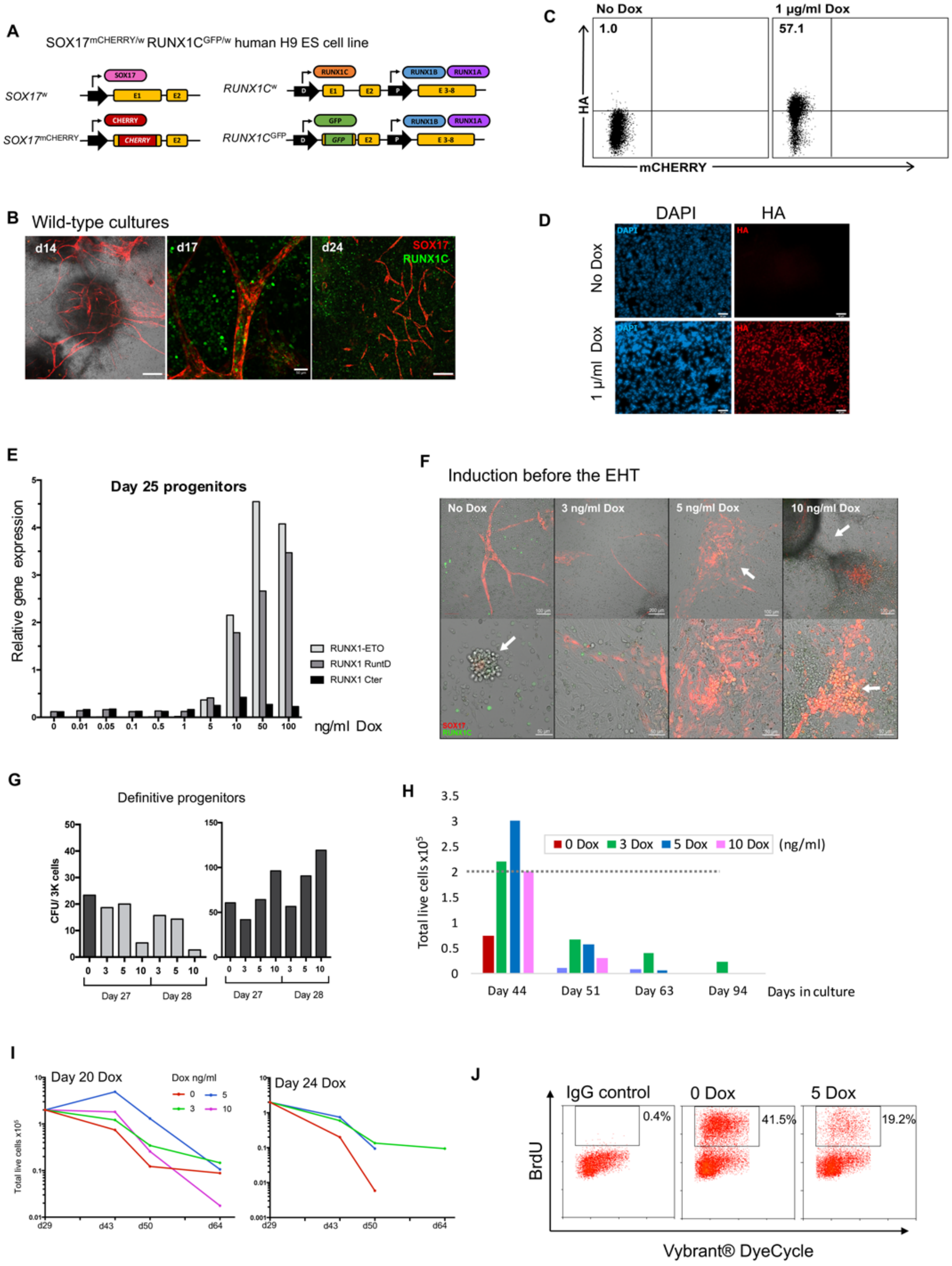

**FIGURE S1. Expression of *RUNX1-ETO* leads to a reversible differentiation and growth arrest of human early hematopoietic progenitor cells**

A) Schematic representation of the targeted alleles in the SOX17$^{mCHERRY/w}$ RUNX1C$^{GFP/w}$ human H9 ES dual reporter cell line. Left: Wild-type and targeted alleles in the *SOX17* locus, with mCHERRY sequence inserted into exon 1. Right: Wild-type and targeted alleles in the *RUNX1* locus. GFP sequence was inserted into exon 1, resulting in expression of GFP from the distal (D) promoter and RUNX1B and RUNX1A isoforms from the proximal (P) promoter within the same allele. Promoters and exons are shown with black arrows and yellow boxes, respectively. Protein products generated from each allele is represented with ovals.

B) RUNX1C+ hematopoietic progenitors emerge from cell clusters located within vascular structures of SOX17+ hemogenic endothelium after the EHT in wild-type EB cultures. Confocal images of EB cultures from the SOX17$^{mCHERRY/w}$ RUNX1C$^{GFP/w}$ human H9 ES double reporter cell line at several days during differentiation, as indicated. SOX17 (mCHERRY) expression marks arterial structures and RUNX1C (GFP) marks hematopoietic progenitors. Scale bar: 200 µm (d14, d24), 50 µm (d17). Fluorescence and brightfield channels are merged at d14 image. At d14: EBs appear as opaque round structures surrounded by a stromal layer containing SOX17+ hemogenic endothelium and arterial structures. At d17: RUNX1C+ blood progenitors are generated from SOX17+ hemogenic endothelium, mimicking structures observed during embryonic AGM hematopoiesis. At d24: progenitors have detached from the endothelium and gone in suspension to further grow and differentiate.

C) Intracellular flow cytometry analysis showing RUNX1-ETO induction upon addition of 1 µg/ml Dox for two days in a pooled population of puromycin-resistant cells after transfection. Detection using an anti-HA tag DyLight® 650-conjugated antibody.

D) Immunofluorescence assay showing RUNX1-ETO induction upon addition of 1 µg/ml Dox for two days in a single-sorted clone. Images are of clone #18 and representative of six single-sorted clones. Cell nuclei are stained with DAPI (blue) and cells expressing HA-*RUNX1-ETO* are detected through an anti-HA antibody (red). Fluorescence channels are merged on right panels. Scale bar: 50 µm.

E) *RUNX1-ETO* and *RUNX1* gene expression in response to Dox titration on d25 hematopoietic progenitors. Primers were designed to amplify: the translocation breakpoint (RUNX1-ETO), the DNA-binding domain present in both gene products (RUNX1 RuntD) and the carboxy-terminal domain present only in endogenous RUNX1 (RUNX1 C-ter). Gene expression was normalized to that of GAPDH.

F) RUNX1-ETO induction before the EHT disrupts the vascular organization and disrupts blood formation. Confocal images of combined Z-stack layers from d16 hematopoietic differentiation cultures with RUNX1-ETO induced from d10 (before the EHT) using 3, 5 or 10 ng/m Dox. Each column shows different fields and magnification within the same Dox-treated culture. White arrows in the 0 Dox condition point emerging RUNX1C-/+ blood progenitors. Arrows in the Dox-treated samples point aberrant structures including disorganized vasculature, RUNX1C- emerging progenitors and *RUNX1C+ SOX17+* co-expressing progenitors. Brightfield and fluorescence field channels are merged. Scale bars: 50, 100 or 200 µm, as indicated. SOX17 (mCHERRY, red) and RUNX1C (GFP, green).

G) RUNX1-ETO causes a reversible differentiation block. Colony-forming unit assays of definitive progenitors from EB cultures treated with Dox during 7 days at day 27 and day 28. For the CFU assays, progenitors were plated in triplicate at a concentration of 3,000 live cells/well in either continued (light grey) or interrupted (dark grey) Dox treatment. Individual graphs correspond to different biological replicates.

H) Previously induced progenitor cells show an initial growth response and increased survival compared to uninduced cells (Related to Figure 2B).

I) Low levels of RUNX1-ETO induction increases the survival of a subset of progenitor cells. Additional examples to what is shown in Figure 2B. Replating assays of hematopoietic progenitors from cultures treated at d20 or d24 with different Dox concentrations, showing two representatives each of three independent experiments. Floating hematopoietic cells were plated at 2 $x10^5$ cells/well in the correspondent Dox concentration and cell numbers were measured weekly at three time points, as indicated. On d24 graph, only 3-dox treated cells were able to survive over 28 days in the replating assays.

J) RUNX1-ETO produces a cell cycle arrest in the G1 phase. Histograms showing cell cycle kinetics of wild-type and RUNX1-ETO-induced progenitor cells. EB cultures were induced at d21 with 5 ng/ml Dox for 4 days and then were pulse-labelled with

25 μM BrdU for 3.5h. Non-adherent cell progenitors were fixed and stained with FITC-conjugated anti-BrdU antibody and Vybrant-DyeCycle. DNA content and cell cycle distribution were analysed by flow cytometric analysis. Boxed cells represent cells that have entered the S-phase of the cell cycle during the BrdU incubation. FITC IgG control is shown. Results correspond a representative of two biological replicates.

Figure S2

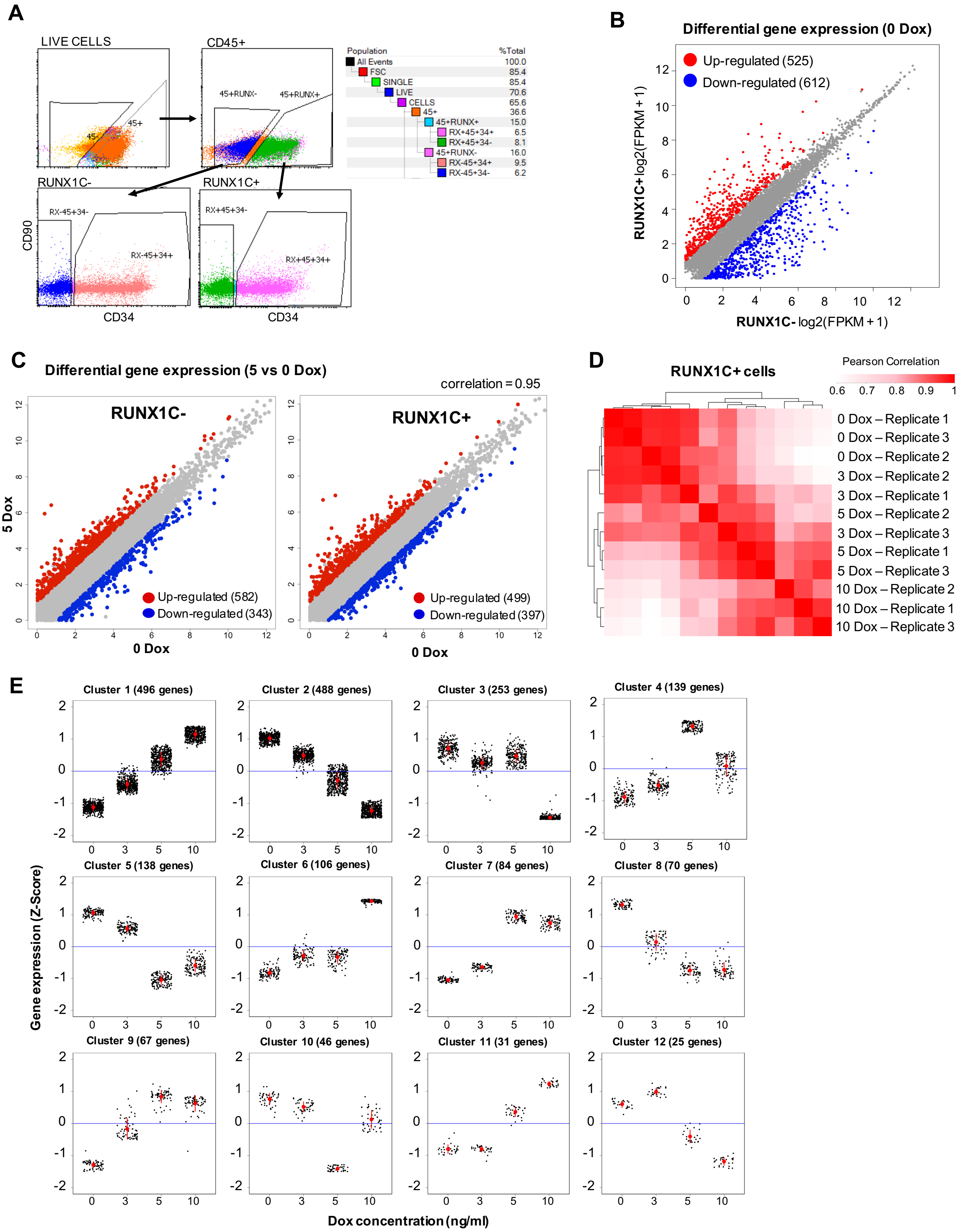

**FIGURE S2: RUNX1-ETO induction leads to cell-type and dose-dependent changes in gene expression**

A) Flow cytometry strategy for sorting of d21 cultures based on CD45 (BV), RUNX1C (GFP/FITC) and CD34 (Pe-Cy7) expression.
B) Clustering of gene expression RNA-Seq data by log2 fold FPKM +1 (fragments per kilobase of transcripts per million mapped reads) values of genes differentially expressed (two-fold change) after RUNX-ETO induction using 5 Dox in both RUNX1C- and RUNX1C+ (CD45+ CD34+) populations. Adjusted P value <0.05.
C) Clustering of gene expression RNA-Seq data by log2 fold FPKM +1 (fragments per kilobase of transcripts per million mapped reads) values of genes differentially expressed (two-fold change) after RUNX-ETO induction using 5 ng/ml Dox in both RUNX1C- and RUNX1C+ (CD45+ CD34+) populations. Adjusted P value <0.05.
D) Hierarchical clustering using Pearson correlation coefficients of gene expression in CD45+ CD34+ RUNX1C+ progenitors upon 24-hour Dox exposure (0, 3, 5 or 10 ng/ml) from three biological replicates.
E) Co-variance analysis of gene expression RNA-Seq data by Z-score from CD45+ CD34+ RUNX1C+ sorted progenitor cells upon RUNX1-ETO induction with 3, 5 or ng/ml Dox for 24h, showing 12 clusters/groups of genes with differential expression response to the level of RUNX1-ETO induction. Number of genes comprised on each cluster are indicated. Black dots represent transcript levels for each individual gene. Red dots and bars represent the mean and standard deviation.

Figure S3

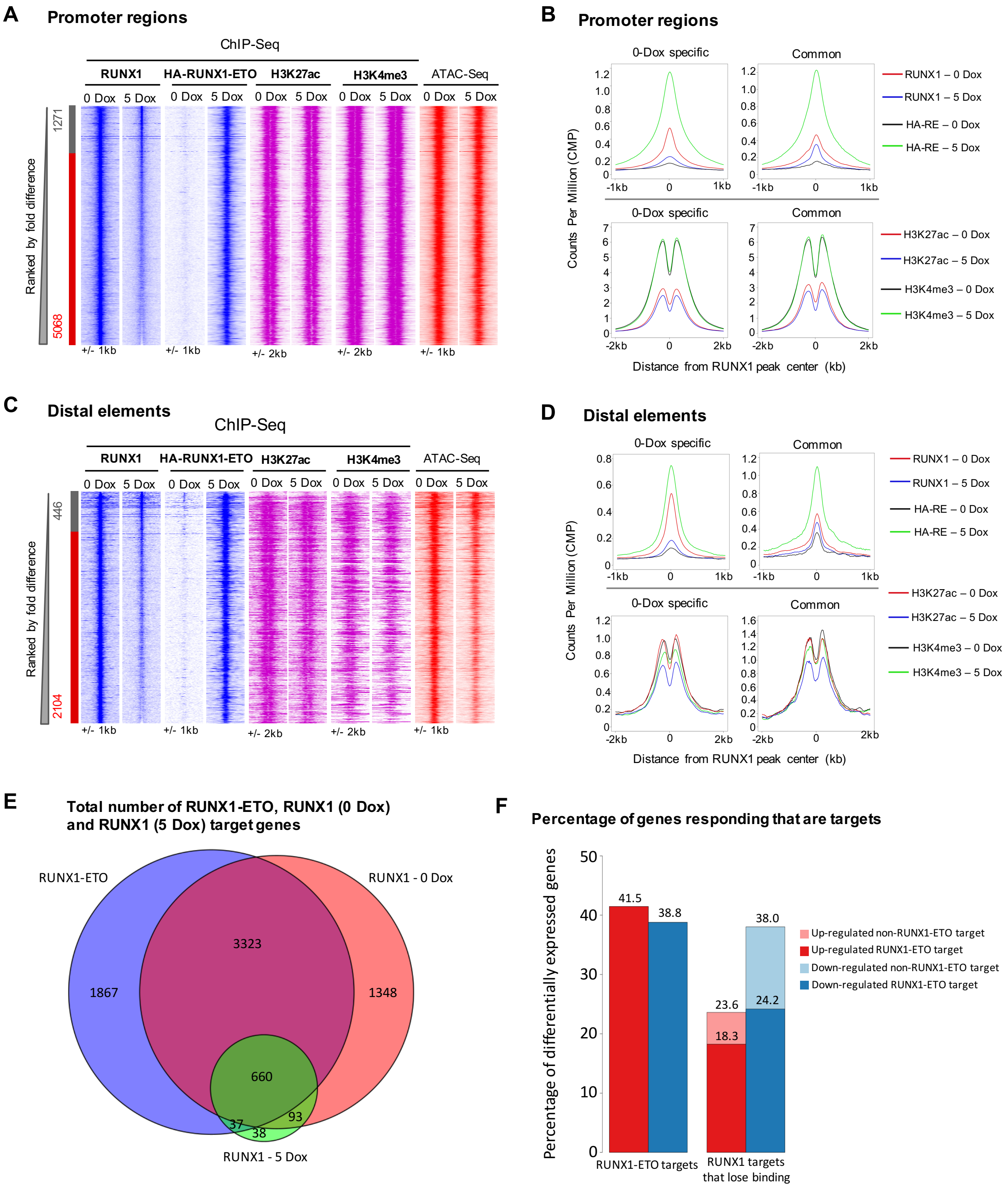

**FIGURE S3: RUNX1-ETO blocks the binding of RUNX1 at distal elements and leads to downregulation of RUNX1 (but not RUNX1-ETO) target genes**

A and C) Comparison of RUNX1 binding at promoter (A) and distal (C) regions from ChIP-Seq in 0 and 5 Dox-treated CD34+ populations ranked by fold difference, considering peaks with enrichments greater than 2-fold between samples to be specific. Sample-specific sites and number of peaks are indicated alongside, being: red the 0-Dox specific and grey the shared peaks. ChIP-Seq enrichment for HA-RUNX1-ETO, H3K27ac and H3K4me3 in each sample and chromatin accessibility peaks are plotted along the same coordinates as the RUNX1 ChIP-Seq promoter (A) and distal element (C) peaks.

B and D) Average profiles for transcription factor (Top panels) and histone modification (Bottom panels) ChIP-Seq data centred on RUNX1 promoter (B) or RUNX1 distal element (D) -binding peaks in the 0 Dox-specific and common peaks.

E) Venn diagram of RUNX1-ETO and RUNX1 ChIP data showing overlap of binding events between the total number of genes targeted by RUNX1-ETO, RUNX1 in 0 Dox uninduced cells and RUNX1 after 5 Dox induction.

F) Graph depicting the percentage of differentially expressed (up- or down-regulated) genes that respond to RUNX1-ETO induction and are either RUNX1-ETO or RUNX1 targets.

Figure S4

A

| Sequencing statistics - 0 Dox | |
|---|---|
| No. of cells sequenced | 3,327 |
| No. of cells passed QC | 3,141 |
| Median no. of reads per cell | 75,452 |
| Median no. of genes per cell | 3,149 |

| Sequencing statistics - 5 Dox | |
|---|---|
| No. of cells sequenced | 3,814 |
| No. of cells passed QC | 3,669 |
| Median no. of reads per cell | 67,098 |
| Median no. of genes per cell | 3,149 |

B

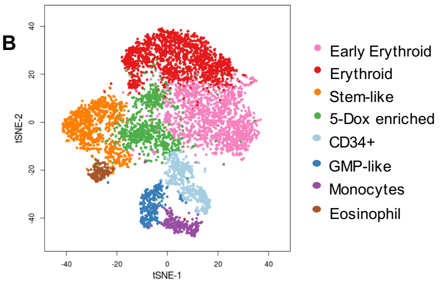

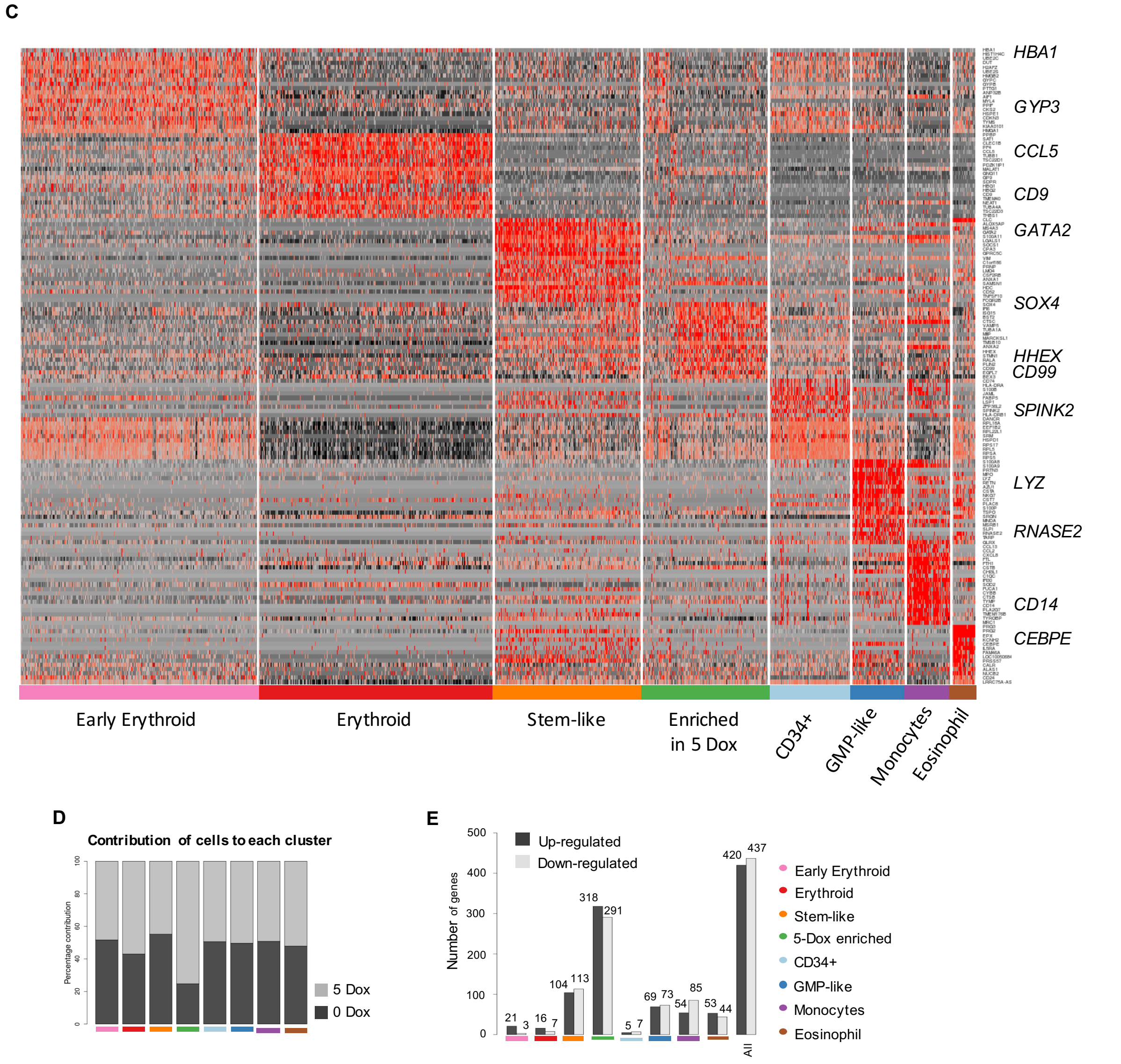

**FIGURE S4: The CD45+CD34+RUNX1C+ population contains precursors from distinct blood lineages as well as multipotent cell progenitors.**

A) Table of the scRNA-Seq sequencing statistics showing the total number of sequenced cells, number of cells that passed the quality control (QC), and the median of reads and sequenced genes per cell for each 0 and 5 ng/ml Dox conditions.

B) Two-dimensional t-SNE maps displaying a total number of 7,135 CD45+ CD34+ RUNX1C+ sorted single cells from the combined data of 0 and 5 Dox treated cells including identified cell populations based on expression of known marker genes.

C) Heatmap showing the expression of the top 20 marker genes specific to each cluster (same colour coding as in (B)). Representative genes from each cluster are indicated.

D) Proportion bars showing the percentage of contribution of 0 and 5 ng/ml Dox dataset to each individual cell cluster.

E) Number of up- and down- regulated genes in each cell cluster upon treatment with 5 ng/ml Dox.

Figure S5

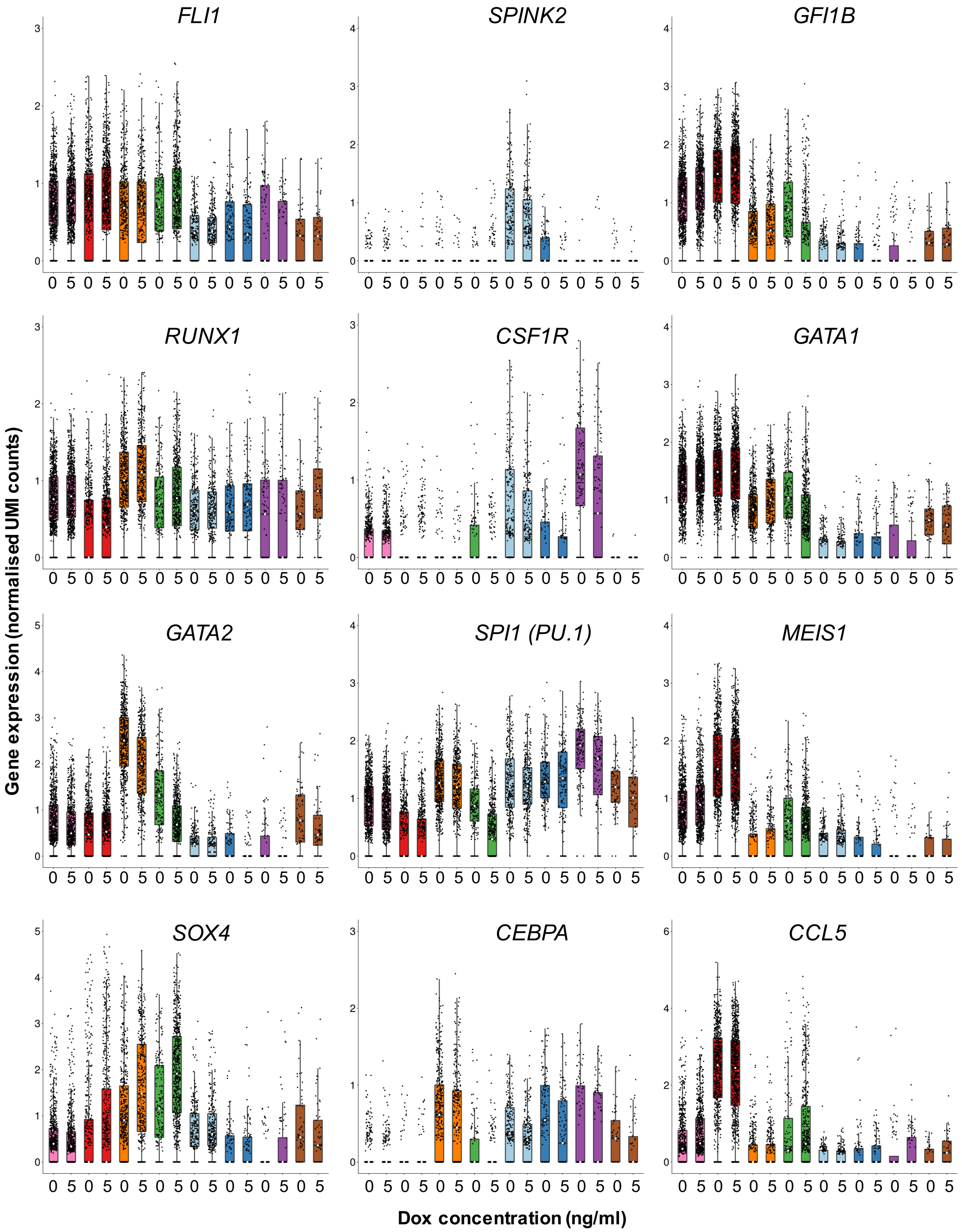

**FIGURE S5: Induction of RUNX1-ETO results in the upregulation of *SOX4* as well as downregulation of important regulators of myelopoiesis in the 5-Dox enriched population**

Box plots indicating expression levels of the individual marker genes in **Error! Reference source not found.** in the different ESC derived cell populations (color coded) and both datasets (0 and 5 Dox). Black dots represent transcript levels in an individual single cell. Boxes and bars represent the mean and standard deviation, respectively.

Figure S6

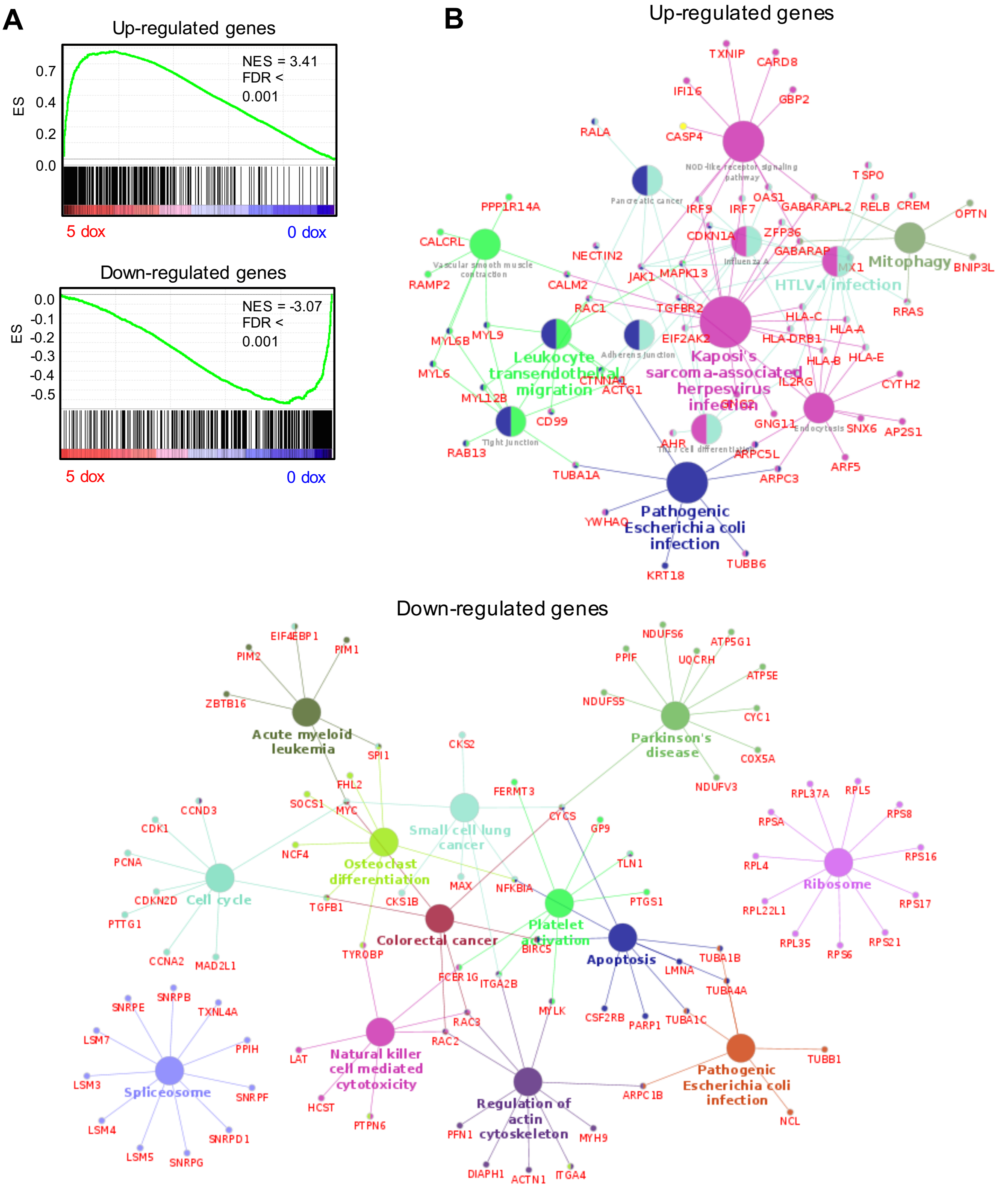

**FIGURE S6: RUNX1-ETO induction yields a similar pattern of transcriptional dysregulation in the 5-Dox enriched and the 5 Dox-treated bulk progenitor populations**

A) Gene Set Enrichment Analysis for correlation of upregulated (top panel) and downregulated (bottom panel) gene signatures between single-cell sorted cells and the CD45+ CD34+ RUNX1C+ bulk populations upon 5-Dox induction for 24 hours. ES, Enrichment Score; NES, Normalized Enrichment Score. FDR, False discovery rate.

B) Network diagram of KEGG pathways for up-regulated (above) and down-regulated (below) genes in the 5-Dox enriched cell cluster of CD45+ CD34+ RUNX1C+ sorted single cells upon 24-hours 5 ng/ml Dox treatment.

#### Supplemental Methods

##### Generation and validation of targeted inducible RUNX1-ETO SOX17$^{mCHERRY/w}$ RUNX1C$^{GFP/w}$ hESC lines

The dual reporter SOX17$^{mCHERRY/w}$RUNX1C$^{GFP/w}$ hESC H9 line was previously generated by us (Ng et al., 2016). RUNX1-ETO cDNA was amplified from the pSIEW-RUNX1-ETO vector with primers including an HA tag sequence following the Kozak sequence containing the start codon and restriction endonuclease sites SalI (5') and MluI (3') for subsequent cloning into the multiple cloning site of the pTREG-CAGGS-Tet3G AAVS1 knock-in plasmid (SUP TABLE1). The pSIEW-RUNX1-ETO and the pTREG-CAGGS-Tet3G vectors were gifts from Olaf Heidenreich (Newcastle, UK) (Bomken et al., 2013) and Su-Chun Zhang (Wisconsin, US) (Qian et al., 2014), respectively. RUNX1-ETO-AAVS1 targeting vectors comprised a 804-bp 5' homology arm, a tetracycline-inducible promoter (TRE3G-CMV) driving expression of HA-RUNX1-ETO cDNA sequence, Puromycin N-acetyltransferase resistance cassette with gene expression to be driven by endogenous promoter after genomic insertion, a CAG promoter driving expression of a modified Tet3G reverse tetracycline-controlled transactivator (rtTA) and a 837-bp 3' homology arm. Vectors were electroporated with a pair of AAVS1 Transcription Activator-Like Effector Nucleases (TALENs) into SOX17$^{mCHERRY/w}$RUNX1C$^{GFP/w}$ hESC H9 cells, which were then selected for Puromycin-resistant colony growth. Single cell sorted clones were screened for transgene insertion by PCR using primers designed to amplify the boundaries of the genomic insertion (SUP TABLE2). Homozygous or heterozygous targeting of the AAVS1 locus was identified with a pair of primers designed to cover both locus sides, resulting in fragment amplification only in presence of a wild type allele. Genomic integrity was confirmed using the Illumina HumanCytoSNP-12 v2.1 array.

##### Maintenance and hematopoietic differentiation of hESC

Culture and enzymatic passaging of hESC lines was conducted as previously reported (Ng et al., 2008). Hematopoietic differentiation of hPSCs was performed following the spin Embryo Body (EB) method in STAPEL medium (Ng et al., 2008) with supplements used as reported (Ng et al., 2016) with some modifications, detailed below. Albumin was substituted for a 1:1 mixture of albumin from rice endosperm (ScienCell Research Labs cat# OsSA) and Bovostar acid stripped Bovine Serum Albumin (BSA, Bovogen cat# BSAS 0.1). For the first 4 days, 7.5

ng/ml ACTIVIN A (ACT, R&D Systems cat# 338-AC) and 0.5 µM CHIR99021 (Tocris Biosciences cat# 4423) were used and 3.5 µM SB431542 (Sapphire Bioscience cat# 13031) and 3 µM CHIR99021 were added 4-6 hours before the day 2 time point and until day 4. After day 4, 5 ng/ml BMP4 and 10 ng/ml Insulin-like growth factor 2 (IGF2, PeproTech cat# 100-12) were used instead of the original concentrations. Adherent cultures after d8 were supplemented with 20 ng/ml BMP4, 100 ng/ml stem cell factor (SCF, PeproTech cat# 300-07), 100 ng/ml FMS-like tyrosine kinase 3 receptor (FLT3) ligand (PeproTech cat# 300-19), 50 ng/ml Thrombopoietin (TPO, PeproTech cat# 300-18), 50 ng/ml vascular endothelial growth factor (VEGF, PeproTech cat# 100-20), 25 ng/ml Interleukin (IL) 3 (PeproTech cat# 200-03), 25 ng/ml IL6 (PeproTech cat# 200-06), 20 ng/ml IGF2, 10 ng/ml basic Fibroblast Growth Factor (FGF2, PeproTech cat# 100-18B) and 1x Penicillin/Streptomycin (Pen/Strep, 5000 U/ml and 5000 U/ml respectively, Thermo Fisher Scientific cat# 15140122). Plates were toped up with media every 3 days and half-media changes were performed when the media capacity of the plate was reached. Upon Dox treatment, adherent cultures were supplemented with a ‘5-factor’ cytokine mix including 100 ng/ml SCF, 100 ng/ml FLT3 ligand, 50 ng/ml TPO, 25 ng/ml IL3 and 25 ng/ml IL-6. For analysis, EBs were harvested at different time points and dissociated into single-cell suspensions using TryPLE select (Invitrogen) for non-adherent EBs and Collagenase Type 1 or Type 4 (Worthington, CLS-1 or CLS-4) for adherent EB cultures and passed through 23- and 25-gauge needles and a 40µm filter (Ng et al., 2008).

**Flow cytometric analysis**

Flow cytometric analysis was performed using BD Fortessa analyser using antibodies against surface antigens detailed in SUPL TABLE 3. For intracellular flow cytometric analysis, cell pellets were fixed with Fixation/Permeabilization solution (BD Pharmingen cat# 554722) on ice for 30 minutes. Cell suspensions were washed with 1x Perm/Wash buffer (BD Pharmingen cat# 554723) and HA-RUNX1-ETO was detected using an primary conjugated antibody aganst the HA-tag (Anti-HA tag [16B12] DyLight® 650-conjugated, Abcam cat# ab117515).

**Cell sorting**

Fluorescence-activated cell sorting (FACS) was done in a FACS Aria cell sorter. Antibodies against CD9 and EpCam were used for sorting undifferentiated hPSCs and against CD34, CD45 and CD90 for sorting hematopoietic progenitors SUPL TABLE 4. Magnetic-activated cell sorting (MACS) of the CD34$^{high}$ hematopoietic cell population was performed using CD34 MicroBeads Kit UltraPure human (Miltenyi Biotec cat# 130-100-453) and a Mini (MS) & Midi (ML) MACS™ Starting Kit (Miltenyi Biotec cat# 130-091-632) as in the manufacturers protocol.

**Intracellular immunostaining**

Adherent cells on 48-well plates were fixed and permeabilized by 15-minute incubation at room temperature with 4% Paraformaldehyde and 0.5% Triton solution and non-specific binding of proteins to the antibody was blocked with 10% FCS Perm/Wash buffer (BD Pharmingen cat# 554723). HA-RUNX1-ETO was detected using an Anti-HA-tag primary conjugated antibody (Anti-HA tag [16B12] DyLight® 650-conjugated, Abcam cat# ab117515) and nuclei were stained with DAPI. Cells were subsequently analysed by epifluorescence imaging.

**Imaging**

Epifluorescence images of the *in vitro* hematopoietic cultures and immunostainings were taken using the 10x and 20x objectives of a Zeiss AxioObserver Z1 microscope and a Zeiss AxioCam monochrome camera and were processed with the Zen Blue software. Confocal images of the *in vitro* hematopoietic cultures were taken with a Zeiss LSM780 microscope using a 10x objective and processed with Zen Black software. All images were exported as separate layers in JPEG format and assembled in Adobe Photoshop when required. Brightness and contrast adjustments were applied equally to all images.

**Colony-forming assays**

Colony-forming-unit (CFU) assays were performed as reported (Ng et al., 2016) with some modifications. Briefly, 3-5 x $10^3$ cells were cultured in 1% methylcellulose, supplemented with the ‘5-factor’ cytokine mix (as used in the hematopoietic differentiation) plus 10 µg/ml human low-density lipoproteins (hLDL, Stem Cell Technologies cat# 02698) and 5 U/ml erythropoietin (EPO, PeproTech cat# 100-64). For the preparation of 1% methylcellulose, 40 ml serum-free 2.6% MethoCult™

H4100 (Stem Cell Technologies #01400) was mixed with an equal volume of 2x STAPEL-P medium (STAPEL medium made with IMDM containing 2x supplements and without PFHMII) plus 20 ml of 1x STAPEL medium to give a final volume of 100 ml. Cells were cultured either with or without Dox and each condition was set up in triplicates in ultra-low attachment 24-well plates (cat# NUN144530). Plates were scored for hematopoietic CFUs after 7 to 10 days.

**Replating assays**

Replating assays were conducted on non-adherent hematopoietic progenitors plated at a known concentration on Matrigel-coated wells of a 6-well plate. Cells were harvested, counted and replated weekly. Live and death cells were determined using a FL Countess-II Automated Cell Counter (Thermo Fisher Scientific) after Tripan-Blue staining.

**Cell cycle analysis**

Cell cycle was analyzed by flow cytometric detection of the incorporation of a thymidine analog (BrdU) and of a fluorescent cell-membrane permeable DNA intercalator. Cells in culture were incubated for 3 hours with 25 µM BrdU (Sigma cat# B5002) and non-adherent progenitor cells were fixed in 75% ethanol. Suspensions of fixed cells were pelleted and re-hydrated with PBS for 20 minutes. Double stranded DNA was denatured by 20-minute incubation with 200 µl 2N HCl, to allow binding of the antibody to the BrdU nucleoside. Cells were washed twice with PBS and washed twice with blocking buffer (5% FBS, 0.1% $NaN_3$, 0.1% TritonC100 in PBS) to bring the pH up to avoid denaturation of the antibodies. Samples were subjected to RNAseA treatment (100 µg/ml) in PBS for 30 minutes at 37°C and IgG control and BrdU stainings were performed using FITC-conjugated antibodies (BD Pharmingen cat# 556028) at room temperature for 50 minutes. Cells were washed with PBS and incubated for 30 minutes with 1 µM Vybrant DyeCycle Violet Stain in PBS prior to flow cytometric analysis.

**Gene expression analysis**

Total RNA was extracted using the Bioline Isolate II RNA Mini Kit according to the manufacturers' instructions. cDNA was reverse-transcribed using random hexamer priming and Tetro cDNA synthesis kit (Bioline) or using Oligo $(dT)_{18}$ priming and SuperScript™ II Reverse Transcriptase (Thermo Fisher Scientific) according to the manufacturers' instructions. Gene expression was analysed by quantitative real-time PCR analysis using Taqman reagents and probes (SUPL TABLE 5) or SYBR Green master mix and primers designed to amplify cDNA fragments (SUPL TABLE 6) (Applied Biosystems). Analyses were performed in technical duplicates and GAPDH was used as the reference gene to normalize data.

**RNA-Seq library preparation**

RNA-sequencing (seq) libraries were prepared using a TruSeq® Stranded mRNA Library Prep (Illumina cat# 20020594) following the Low Sample (LS) workflow according to manufacturer's instructions. Libraries were subjected to a quality control using a High Sensitivity DNA chip on an Agilent Technologies 2100 Bioanalyser™ instrument and were quantified using the RT-qPCR-based method KAPA Library Quantification Kit for Illumina Sequencing Platforms (Roche cat# KR0405), following the manufacturer's protocol. Libraries were run in a pool of twelve indexed libraries in a NextSeq (Illumina) machine using sequencing by synthesis chemistry and a NextSeq® 500/550 High Output 150 cycle sequencing kit v2 (cat# FC-404-2002), obtaining 75 bp paired-end reads.

**Assay for transposase-accessible chromatin with high-throughput sequencing (ATAC-seq)**

Chromatin accessibility was evaluated by ATAC-seq using a modified protocol to as reported (Buenrostro et al., 2015; Corces et al., 2016). Briefly, 50,000 single-cell sorted hematopoietic progenitors were pelleted and snap-frozen upon resuspension in 5 µl sucrose freezing buffer, consisting of 60 mM KCl, 15 mM NaCl, 5 mM $MgCl_2$, 10 mM Tris pH 7.4 and 1.5 M sucrose. Transposition reaction was performed for 30 minutes at 37°C upon addition of 45 µl of ATAC reaction mix consisting of 25 µl Tagmentation DNA Buffer (Illumina cat# FC-121-1030, Nextera DNA Library Prep Kit), 2.5 µl Tn5 Transposase enzyme (Illumina cat# FC-121-1030, Nextera DNA Library Prep Kit), 1 µl of 0.5% Digitonin (Promega cat# G9441) and 16.5 µl water. DNA was purified using a MinElute Reaction Cleanup Kit (Qiagen cat# 28204). Purified DNA fragments were amplified using Customized Nextera PCR Primer Adaptors and NEBNext High-Fidelity 2x PCR Master Mix (New England Biolabs cat# M0541). Optimal number of cycles, prior reaching saturation of the PCR in order to reduce GC and size bias, was determined by monitoring the reaction as previously described (Buenrostro et al., 2015). Adaptor dimers were cleaned up from the libraries uwing AMPure magnetic beads (Beckman Coulter) prior to validation. Libraries were evaluated using a High Sensitivity DNA chip on an Agilent Technologies 2100 Bioanalyser™ instrument and concentration was measured using a RT-qPCR-based method (KAPA Library Quantification Kit for Illumina Sequencing Platforms). Libraries were also validated by RT-qPCR evaluation of the ratio of open (TBP promoter) to closed regions of DNA (chromosome 18) and active gene body (β-actin). Libraries were sequenced in a pool of twelve indexed libraries in a NextSeq (Illumina) machine and a NextSeq® 500/550 High Output 75 cycle sequencing kit v2 (cat# FC-404-2005), obtaining 75 bp single-end reads at the Genomics Birmingham sequencing facility.

**Chromatin Immunoprecipitation with high-throughput sequencing (ChIP-Seq)**

ChIP was performed on CD34+ magnetic sorted samples (RUNX1-ETO and RUNX) and on non-adhered floating mixed progenitors (H3K4me3 and H3K27ac) after single crosslink, as reported (Obier et al., 2016). Antibodies were used against the following antigens (manufacturer, cat #): HA tag (Sigma, H6908), RUNX1 (ab23980, Abcam), H3K27ac (Abcam, ab4729), H3K4me3 (,), H3K79me2 (Abcam ab3594). For H3K79me2 ChIP, a concentration of at least 0.2% SDS on the chromatin suspension was used and immunoprecipitation was carried overnight. ChIP libraries for Illumina sequencing were prepared using the KAPA Hyper Prep Kit (Roche, KR0961), as detailed by the manufacturer. Quality control of the libraries was performed using a High Sensitivity DNA chip on an Agilent Technologies 2100 Bioanalyser™ instrument and libraries were quantified using the RT-qPCR-based method KAPA Library Quantification Kit for Illumina Sequencing Platforms, following the manufacturer's protocol. Libraries were sequenced in a pool of twelve indexed libraries in a NextSeq (Illumina) machine and a NextSeq® 500/550 High Output 75 cycle sequencing kit v2 (cat# FC-404-2005).

**Single Cell RNA-Seq (scRNA-Seq)**

Non-adherent progenitors at day 22 of differentiation (untreated and 24-hour Dox treatment) were sorted for CD45+ CD34+ and RUNX1C+. Cells were re-suspended in 80 µl at a concentration of 100-1200 cells/µl for evaluation of cell viability prior to loading of 4000 single cells on a Chromium Single Cell Instrument (10X Genomics).

Library generation for scRNA-seq was performed by the Genomics Birmingham Sequencing Facility using the Chromium Single Cell 3′ Library and Gel Bead Kit v2 (10X Genomics). Libraries were paired-end sequenced on an Illumina NextSeq machine using the cycle parameters recommended by 10X Genomics.

**Statistical analysis**

Experiments were analysed using GraphPad Prism versions 5–7 (GraphPad Software Inc.) and Microsoft Excel (Microsoft corporation).

**Bioinformatic Data processing and analysis**

**Bulk RNA-Seq data analysis**

Sequencing adaptors and low quality bases were trimmed from the raw RNA-Seq reads using Trimmomatic v0.32 (Bolger et al., 2014). The processed reads were then aligned to the human genome (version hg38) using Hisat2 v2.1.0 (Kim et al., 2015) with default settings. Gene expression was measured as fragments per kilobase of transcript per million mapped reads (FPKM) values using with Stringtie v.1.3.3 (Pertea et al., 2015) with default settings. Gene models from the RefSeq database (O'Leary et al., 2016) were used as the reference transcriptome. Only genes that were expressed with an FPKM > 1 in at least one of the samples were retained for further analysis. The raw FPKM values were quantile normalized using the Limma package v3.26.9 (Ritchie et al., 2015) in R v3.5.1. The normalized data was then log2-transformed, with a pseudocount of 1 being added to each of the FPKM values prior to transformation.

Differential gene expression analysis was carried out using Limma. A gene was considered to be differentially expressed if it had a greater than 2-fold change between experimental conditions, and a Benjamini-Hochberg adjusted p-value < 0.05. Kyoto encyclopedia for genes and genomes (KEGG) pathway enrichment analysis was done using the ClueGO plugin v2.5.0 (Bindea et al., 2009) for Cytoscape v.3.61 (Shannon et al., 2003). This was done using a right-sided hypergeometric test, with Benjamini-Hochberg p-value correction for multiple testing. A pathway was deemed to be significantly enriched if the adjusted p-value was < 0.05.

Hierarchical clustering of RNA-Seq samples and replicates was done by first calculating the Pearson correlation value for each pair of samples. The resulting correlation matrix was then hierarchically clustered using complete linkage clustering of the Euclidean distances, and finally plotted as a heatmap in R.

To carry out gene expression co-variance analysis, gene expression values were first transformed to Z-scores using the scale function in R. These were then hierarchically clustered using complete linkage of the Euclidean distances. Clusters corresponding to sets of genes with similar patterns of expression were then extracted from the dendrogram using the dynamicTreeCut package v1.63 (Langfelder et al., 2008) in R using the hybrid method with a minimum cluster size of 25 genes.

To compare the gene expression profile of the RUNX1-ETO induced cells to that of AML patients with the t(8;21) translocation, RNA-Seq data from t(8;21) patients and from healthy peripheral blood stem cells (PBSCs) from *Assi et al*., 2019 was downloaded from GEO using the accession GSE108316. These data were aligned and processed as described above. The sets of genes that were up and down regulated in the RUNX1-ETO induced cells was then compared to the gene expression profiles of the t(8;21) AML cells and PBSCs using gene set enrichment analysis (GSEA) using the GSEA software (Subramanian et al., 2005).

**ATAC-Seq data analysis**

Single-end reads from ATAC-Seq experiments were processed to remove low-quality bases and sequencing adaptors using Trimmomatic. Reads were then aligned to the human genome (version hg38) using Bowtie2 v2.2.6 (Langmead and Salzberg, 2012) with the parameter --very-sensitive-local. Reads that aligned to the mitochondrial genome were removed from further analysis. Potential PCR duplicated reads were identified and removed from the alignments using Picard v2.10.5 (http://broadinstitute.github.io/picard). Open chromatin regions (peaks) were identified using MACS2 v2.1.1 (Zhang et al., 2008) using the settings --nomodel --nolambda -B --trackline. The resulting peaks were then filtered against the hg38 blacklist and simple repeat tracks from the UCSC table browser (Karolchik et al., 2004) to remove any potential artifacts. Peaks were annotated to the nearest gene, and then further annotated as either a promoter of distal element using the annoatePeaks.pl function in the Homer software package v4.9.1 (Heinz et al., 2010).

A peak was annotated as being within a gene promoter if it was within 1.5kb of a transcription start site (TSS) and as a distal element otherwise.

ATAC peak unions were constructed by merging peaks that had summit positions within 400bp of each other. In these cases, peaks were combined to a single peak with a new summit position defined as the mid-point between the summit positions of the original peaks. These average peak positions were used in all further downstream analysis.

To identify regions of differential chromatin accessibility, a peak union was first created for each pair of samples being considered. The read density for these peaks was then retrieved directly from the bedGraph files produced by MACS2 using the annotatePeaks.pl function in Homer with the parameter -size 200. These tag counts were normalized as counts per million (CPM) in R, and further log2-transformed with a pseudocount of 1 added to each value prior to transformation. A peak was considered to be differentially accessible if the fold-difference of the normalized tag count was greater then 2 between experiments. Motif enrichment analysis was then carried out in these sets of peaks using the findMotifsGenome.pl function in Homer.

To create read density plots, peaks were first ordered according to fold-difference. The read density in a 2kb window centered on the peak summits was then calculated using from the bedGraph files produced by Homer using the annotatePeaks.pl file in Homer, using the options -size 2000 -hist 10 -ghist. These were then plotted as heatmaps using java TreeView v1.1 (Saldanha, 2004).

ATAC-Seq data from hematopoietic cell type in various stages of differentiation were obtained from *Corces et al*., 2016 via GEO using the accession GSE74912. These data were aligned and processed as described above.

**ChIP-Seq data analysis**

Reads from ChIP-Seq experiments were processed, aligned to the human genome and de-duplicated in the same way described above for the ATAC-Seq data. Peaks from ChIP-Seq experiments targeting the transcription factors RUNX1 and RUNX1-ETO were identified using MACS2 with default settings. These peaks were then compared to the ATAC-Seq data, with only peaks that occurred within open chromatin regions being retained for further analysis. To identify differential binding of RUNX1 between the 0 and 5 Dox datasets, a union of RUNX1 peaks was first constructed by merging peaks that had summits within 100bp of each other. The read density in these peaks was then retrieved using the annoatePeaks.pl function in Homer and normalized as counts per million in R. Peaks that had a fold-difference of at least 2 were considered to be differentially bound between experiments. RUNX1 and RUNX1-ETO target genes were identified by annotating each peak to its closest TSS using the annotatePeaks.pl function in Homer.

Peaks corresponding to the histone modifications H3K27ac and H3K4me3 were called using MACS2 with default settings. These peaks were then filtered against the hg38 blacklist and simple repeat tracks from the UCSC table browser to remove any potential artifacts.

**Construction of average profiles**

Average profiles for ATAC and ChIP-Seq data were constructed by first normalizing each of the alignment tracks as counts per million (CPM) using the bamCoverage function in deepTools v3.2.0 (Ramírez et al., 2016). These were then plotted using the plotProfile function in deepTools.

**Single cell RNA-Seq data analysis**

Illumina base call (BCL) files that were generated using the Chromium platform from 10x genomics were de-multiplexed and converted to the fastq format using the mkfastq function in CellRanger v2.1.1. These were then aligned to the human genome (version hg38) using the count function in CellRanger. Gene models from the RefSeq database were used as the reference transcriptome. Unique molecular identifier (UMI) counts were processed and normalized using the Seurat v2.3.4 package (Butler et al., 2018) in R. Cells with less than 1500 detectable genes, or that had more than 10% of UMIs aligned to mitochondrial genes were removed from further analysis. Additionally, genes that were detected in less than 20 cells were also excluded from analysis. The cell cycle stage for each cell was inferred using the CellCycleScoring function in Seurat. The possible effects of cell cycle stage, as well as sequencing depth (as measured by the total number of UMIs) per cell were removed from the analysis by linear regression using the ScaleData function in Seurat.

Clustering of cells was performed by first combining the datasets from the 0 and 5 dox treated cells into a single dataset using canonical correlation analysis (CCA). This combined dataset was then clustered using the t-distributed stochastic neighbor embedding (t-SNE) method. Cell clusters were identified using the FindClusters function in Seurat, using a resolution value of 0.4. Cell marker genes, corresponding to genes that are enriched on one cluster relative to others, were identified using the FindMarkers function. A gene was considered as a marker gene if it had a log fold-change value greater than 0.5 and could be detected in at least 50% of cells in that cluster. Differential gene expression analysis was also carried out for each cluster using the FindMarkers function, with genes with a log-fold-change greater than 0.25 and an FDR < 0.05 being considered to be differentially expressed.

Cell trajectory (pseudotime) analysis was carried out using Monocle v2.10.1 (Qiu et al., 2017; Trapnell et al., 2014). Normalized UMI counts from Seurat were first imported into Monocle using the importCDS function. Cells were then ordered along a pseudotime trajectory using the discriminative dimensionality reduction with trees (DDRTree) method using the complete set of cell marker genes identified by Seurat to order the cells.

#### Supplementary tables

##### Table S1: Primers used for cloning

| Oligonucleotide | Sequence | Orientation | Description |
|---|---|---|---|
| SalI_HA-RUNX1 | TACCGTCGACCCGCCATGTACCCATACGACGTCC CAGACTACGCTCGTATCCCCGTAGATGCCAGCAC GA | Forward | RUNX1/ETO cloning into AAVS1 plasmid |
| ETO_MluI | CGCAACGCGTCTACTAGCGAGGGGTTGTCTCTA | Reverse | RUNX1/ETO cloning into AAVS1 plasmid |

##### Table S2: Primers for PCR amplification of genomic DNA in transgene screening assays

| Oligo name (binding) | Orientation | Sequence |
|---|---|---|
| AAVS1 5' screen (AAVS1 5') | Forward | GGACCACTTTGAGCTCTACT |
| AAVS1 5' screen (T2A) | Reverse | TCCACGTCACCGCATGTTAG |
| AAVS1 3' screen (TET3G) | Forward | TGCCTGCTGACGCTCTTGACGATT |
| AAVS1 3' screen (AAVS1 3') | Reverse | GAAGGATGCAGGACGAGAAA |
| Wild-Type screen (AAVS1 5') | Forward | CCCCTATGTCCACTTCAGGA |
| Wild-Type screen (AAVS1 3') | Reverse | CAGCTCAGGTTCTGGGAGAG |

##### Table S3: Conjugated antibodies used for flow cytometry

| Antibody | Fluorochrome | Manufacturer | Catalogue # |
|---|---|---|---|
| CD16 | PeCy7 | BioLegend | 302015/302016 |
| CD34 | BV-421 | BioLegend | 343609 |
| CD34 | PeCy7 | BioLegend | 343516 |
| CD38 | APC | BD Pharmingen | 555462 |
| CD45 | BV-421 | BioLegend | 304032 |
| CD90 | APC | BD Pharmingen | 559869 |
| CD90 | BV-421 | BioLegend | 328121 |

##### Table S4: Conjugated antibodies used for single cell sorting

| Antibody | Fluorochrome | Manufacturer | Catalogue # | Stock | Dilution |
|---|---|---|---|---|---|
| CD9 | PE | BD Pharmingen | 555372 | 100 tests | 1:100 |
| CD34 | PeCy7 | BioLegend | 343516 | 100 µg/ml | 1:100 |
| CD45 | BV-421 | BioLegend | 304032 | 25 µg/ml | 1:50 |
| CD90 | APC | BD Pharmingen | 559869 | 0.2 mg/ml | 1:100 |
| CD326 (EpCam) | BV-421 | BioLegend | 324220 | 100 tests | 1:30 |

#### Table S5: Taqman probes used for gene expression analysis from total RNA

| Name | Probe |
|---|---|
| GAPDH | Hs99999905_m1 |
| GATA1 | HS00231112_m1 |
| GFI1 | Hs00382207_m1 |
| GFI1B | Hs01062469_m1 |
| PU.1 | HS00231368_m1 |
| RUNX1 COMMON | HS00231079_m1 |
| RUNX1C | Hs01021967_m1 |
| RUNX1T1 | Hs00231702_m1 |

#### Table S6: Primers used for gene expression analysis from total RNA

| Primers for cDNA | Forward | Reverse |
|---|---|---|
| *GAPDH* | CCTGGCCAAGGTCATCCAT | AGGGGCCATCCACAGTCTT |
| *RUNX1 (Cter)* | CCCTCAGCCTCAGAGTCAGAT | AGGCAATGGATCCCAGGTAT |
| *RUNX1 (Runt Domain)* | AACAAGACCCTGCCCATCGCTTTC | CATCACAGTGACCAGAGTGCCAT |
| *RUNX1/ETO junction* | TCAAAATCACAGTGGATGGGC | CAGCCTAGATTGCGTCTTCACA |
| DHFR-KRAS | AATTCCACGATGCTGATGCG | CAAGGCACTCTTGCCTACGC |
| KIT-DHFR | TCAATTCTGTCGGCAGCACC | CATGGCGTTTTCCATGCCGA |
| *RUNX1/EVI1 junction* | CCACAGAGCCATCAAAATCA | TCTGGCATTTCTTCCAAAGG |
| *EVI1 exon 7* | AAACCTTTGCCGTCATAAGCG | CGTAGTGCTGAACATTTGTCCACAG |
